# Physics-inspired accuracy estimator for model-docked ligand complexes

**DOI:** 10.1101/2024.10.17.618358

**Authors:** Byung-hyun Bae, Jungyoon Choi, Chaok Seok, Hahnbeom Park

## Abstract

Model docking, which refers to ligand docking into the protein model structures, is becoming a promising avenue in drug discovery with the advances in artificial intelligence (AI)-based protein structure prediction. However, a significant challenge remains; even when sampling was successful in model docking, typical docking score functions fail to identify correct solutions for two-thirds of them. This discrepancy between scoring and sampling majorly arises because these scoring functions poorly tolerate minor structural inaccuracies. In this work, we propose a deep neural network named DENOISer to address the scoring challenge in model-docking scenario. In the network, ligand poses are ranked by the consensus score of two independent sub-networks: the *Native-likeness prediction* and the *Binding energy prediction networks*. Both networks incorporate physical knowledge as inductive bias in order to enhance pose discrimination power while ensuring tolerance to small interfacial structural noises. Combined with Rosetta GALigandDock sampling, DENOISer outperformed existing docking tools on the PoseBusters model-docking benchmark set as well as on a broad cross-docking benchmark set. Further analyses reveal the physics-based components and the consensus ranking approach are the two most crucial factors contributing to its ranking success. We expect DENOISer may assist future drug discovery endeavors by providing more accurate structural models for protein-ligand complexes. The network is freely available at https://github.com/HParklab/DENOISer.

## 1. Introduction

Understanding interactions between proteins and ligands is an essential part of structure-based drug design. One of crucial methods for investigating these interactions is molecular docking, in which energetically most plausible ligand binding conformations are predicted within the receptor protein context. This computational approach allows researchers to explore the ligand binding mode and affinity on target proteins to guide rational design of novel drug candidates.

Docking scenarios can be categorized into three classes depending on the type of information available for receptors of interest. *Self-docking (or re-docking)* involves docking a ligand back into its own binding site from the original bound-form protein structure, which is less realistic yet broadly accepted for benchmarking purposes ^1,2^. *Cross-docking* involves docking a ligand into the same protein but at a different conformation (i.e. bound to other ligands or unbound), and corresponds to a realistic scenario frequently occurring in structure-based drug design ^3^. In the third category, *model docking*, which is the main focus of this work, predicted protein structures replace experimental structures at which ligands are docked into ^4^.

The concept of model docking has gained large attention in recent years. Significant advancements in deep learning methods within biological domains have led to breakthroughs in the protein structure prediction. This breakthrough led to a large increase in the coverage of druggable protein receptors whose structural quality at binding sites approach that of cross-docking receptors. If we set “predicted model confidence of > 0.9 by AlphaFold2” as a criterion for a druggable receptor, nearly 30% of human proteins can be additionally targeted for drug discovery via model docking, when only 15-20% were druggable using experimental structures ^5^.

However, it is more of the prediction accuracy of a docking tool which is hindering the utility of model docking than the quality of predicted receptor models ^4^. The major reason is due to a common pathology shared by score functions in docking tools. Because the majority of docking score functions are tailored to best perform in self-docking scenarios, they lack robustness against noise at the backbone or side-chain level in more realistic cross- or modeling-docking scenarios (**Figure 1a**), leading to selection failures to two-thirds of the cases even sampling was successful (**Figure 1b**). Deep learning based score functions are suggested to remedy these issues, but their generalizability to novel targets are questioned, majorly due to the sparsity of protein-ligand complex data compared to the number of parameters in those networks ^6–10^. Consequently, there is a pressing need for deep learned scoring functions robust not only to model-docking scenarios but also generalizable to novel ligands or receptors.

**Figure 1.**
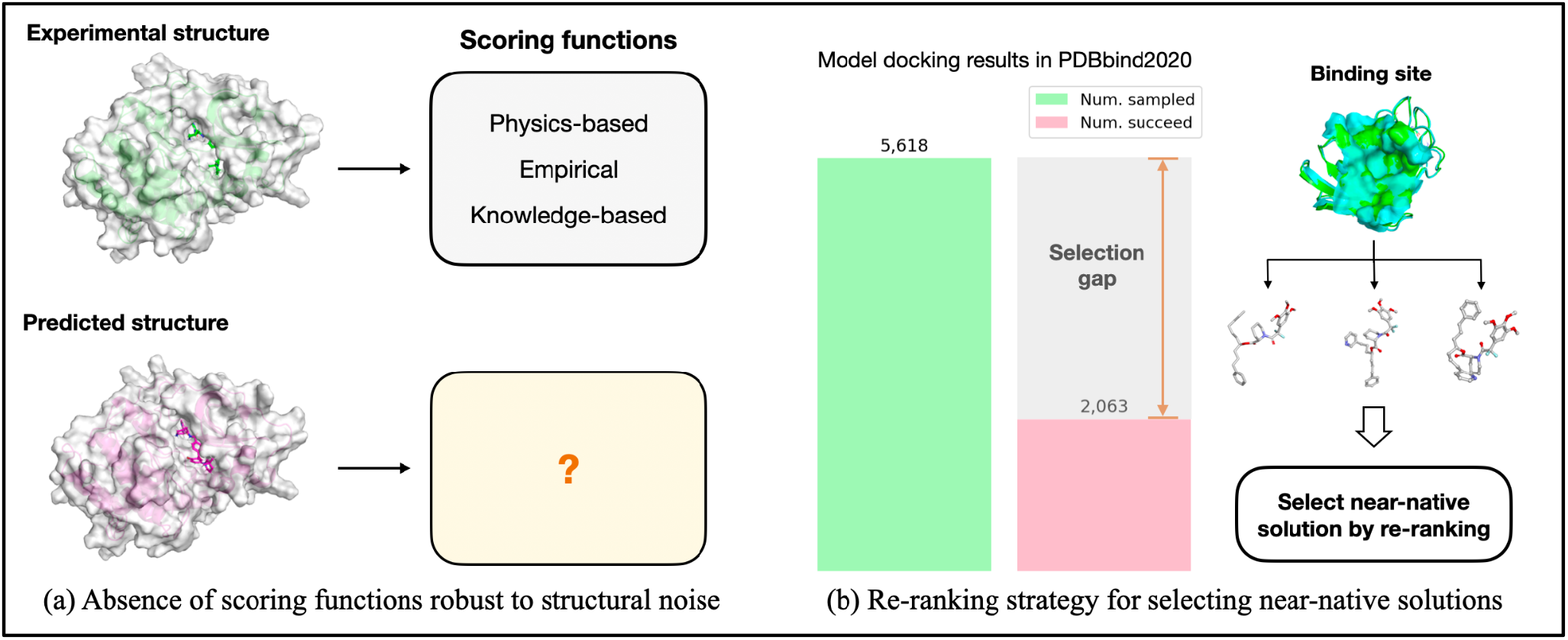
The background and motivation of our method. (a) The protein structure below (magenta) is predicted by the protein structure prediction model (e.g. AlphaFold2, RosettaFold, etc). (b) Strategy for selecting among sampled structures even if the binding site is slightly different. The model docking was benchmarked on the curated 13,644 PDBbind 2020 data set ^11^ using AlphaFold2 for receptor modeling.

We reasoned physical principles can play a key role as inductive bias to a deep learning network to achieve both noise toleration and generalization. Instead of applying exact force field terms, which are often too sensitive to steric clashes and require perfect van der Waals interactions ^12^, the network is designed to put more weights on evaluating globular physical plausibility (e.g. solvation, electrostatic compatibility) at protein-ligand interfaces. With this strategy, a large number of parameters can be regularized to better explain physical observations, and hence can be robust to atomic noises and less prone to overfit to train data alone.

With these considerations, here we present DENOISer (Docking Evaluation for Non-Optimal Interfacial Structures). Our contributions can be summarized as follows. Firstly, we introduce a deep learning-based evaluator robust at structural uncertainties inherent in predicted or cross-docking receptor structures. This approach offers a more realistic solution to docking problems, particularly when dealing with novel protein targets lacking experimental structures. Secondly, we demonstrate that the incorporation of physics-inspired knowledge into the model architecture can considerably enhance its learning efficiency and generalizability. Thirdly, we demonstrate the feasibility of the re-ranking strategy not only for model docking but also for docking to unbound form (apo) proteins, most challenging cross-docking problems, exhibiting structural disparities between the apo and holo forms.

## 2. Method

### 3.1. Overview

As illustrated in **Figure 2**, DENOISer takes multiple poses from docking methods and subsequently processes these input-featured files to generate graph representations of protein-ligand complexes. The resulting graph is then fed into two sub-networks: the *Native-likeness prediction network* and the *Binding energy prediction network*. The final score is obtained by averaging min-max normalized scores each ranging from 0 to 1. Networks are commonly trained using modeled docked decoys for curated 18,094 protein-ligand complexes deposited in PDBbind v.2020, with each complex containing 101 decoys. Receptors are generated using AlphaFold2 (AF2) ^13^, and ligands are docked using Rosetta GALigandDock (GALD) with receptor flexibility ^14^. Further details regarding graph construction and network architectures are described in the following sections.

**Figure 2.**
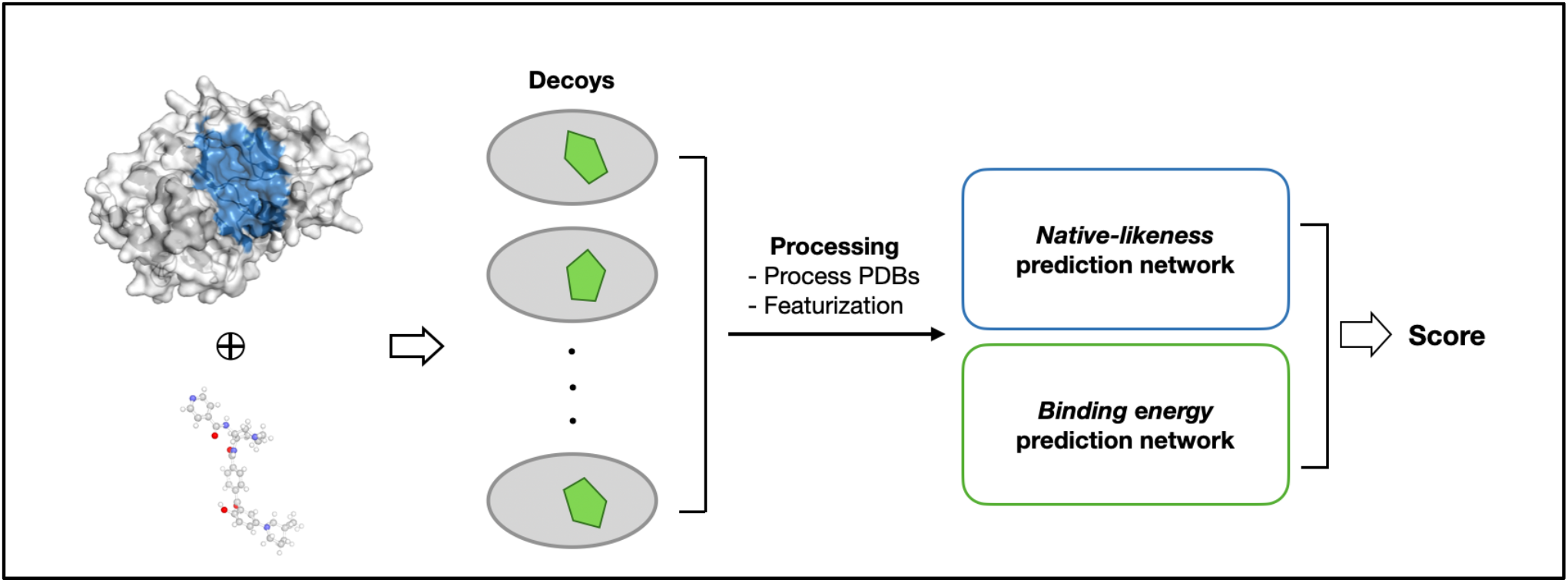
Overview of DENOISer. The gray circles and green pentagons in decoys represent proteins and ligands, respectively. These complexes have the same protein and ligand pair but have different conformations. The final score is obtained as an ensemble of the results of the two networks.

### 3.2 Native-likeness prediction network

#### 3.2.1 Overall model architecture

Three primary types of input graphs are used by the networks: residue graph, atom graph, and local atom graph (**Figure 3a**, details in the next section). All graphs are processed through the graph embedding module (**Figure 3c**) to compute i) the 1-D embedding: *node-wise embedding vectors* through an MLP module and ii) the 2-D embedding: *physics-inspired interaction difference matrices* through an E(n) Equivariant Graph Neural Network (EGNN) ^15^ module. At the 1-D track, node-wise embeddings are learned which conceptually contain ligand-atomwise native-to-decoy structural difference information. At the 2-D track, pairwise interactions are evaluated: global embedding surrounding the binding pocket learned at residue-level is fed along with additional atomic features into atom graphs, then processed through individual EGNN modules. The output node-wise embeddings of protein and ligand are then pairwise concatenated and transformed into 2-D *physics-inspired interaction difference matrices* where structural errors are estimated from four physical types of protein-ligand interactions. Finally, 1-D and 2-D embedding are processed together to estimate two Local Distance Difference Test (lDDT) ^16^ values: *complex lDDT* and *atomic lDDT*. More details about the graph construction, physics-inspired interaction difference matrices, and definitions of two lDDT values are elaborated individually below.

**Figure 3.**
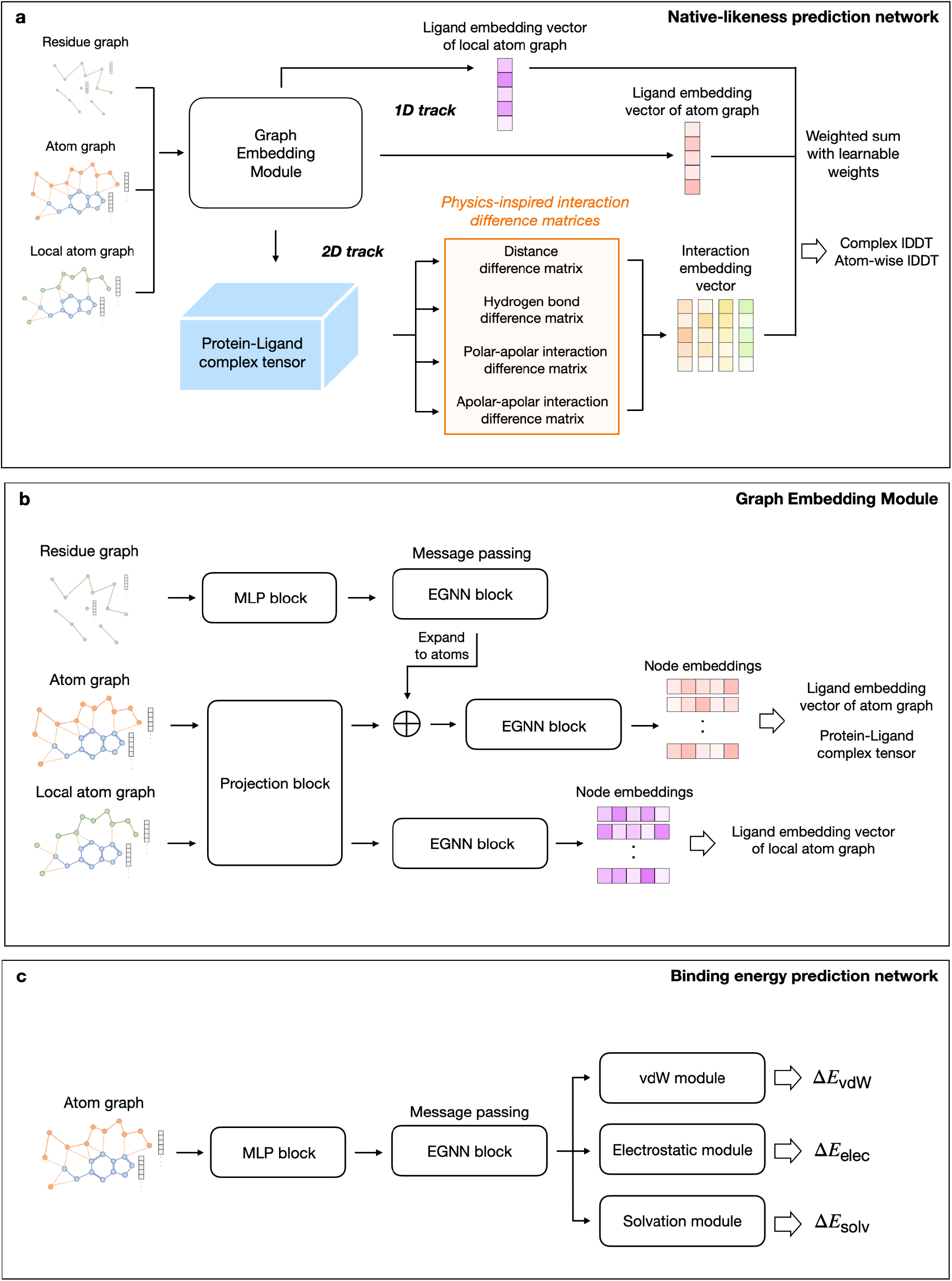
Network architectures of the Native-likeness prediction network and the Binding energy prediction network. (a) Native-likeness prediction network. (b) Architectural details of the graph embedding module in the Native-likeness prediction networks. (c) Energy prediction network. In both panels, vdW denotes van der Waals interaction and EGNN denotes E(n) equivariant graph neural network.

#### 3.2.2 Input graphs

The residue graph operates at a residue-level, in which nodes represent the alpha carbons of protein residues and the ligand atom closest to its center of mass. The atom graph operates at the atom-level resolution, where all atoms except for non-polar hydrogen are expressed as nodes. This graph is constructed with atoms within a 9 Å radius from any atom in the ligand (otherwise excluded from the graph for efficiency). The initial features of nodes and edges for all graphs are reported in **Table S1**. Each graph incorporates edges denoting chemical bonds and spatial connections, where spatial edges are connected if the Euclidean distance between nodes falls below pre-defined cutoffs: 8.0Å cutoff for the residue graph, 4.5Å for the atom graph, and 3.0Å for the local atom graph, respectively. The local atom graph shares the same resolution as the atom graph but is introduced to catch direct intermolecular contacts only. This graph enables more sensitive discerning of near-native structures but is more prone to overfitting, therefore its contribution to the prediction is fixed to 0.1 (out of total 1.0).

#### 3.2.3 Physics-inspired interaction difference matrices

Four types of physical interactions are considered while designing the 2-D track of the network: 1) raw atomic pair distances, 2) hydrogen bonding (i.e. favorable polar-polar interactions), 3) polar-apolar interactions, and 4) apolar-apolar Interactions. From these four interaction difference matrices (orange box in **Figure 3**), per-term interaction differences of the given input over the experimental structure are learned. The major objective of calculating these matrices is to guide lDDT prediction from more *physically regularized and explainable* 2-D embeddings; to this aim auxiliary losses are introduced to tether these 2-D predictions to known labels. We also reasoned that these all-by-all atomic pair predictions may help the network infer structural differences originating from long range effects that are rather diluted in the graph-level learning. Brief descriptions of per-term formulations and auxiliary losses resulting from those matrices are described below (details in **Supplementary Methods**).

##### Distance difference loss

The distance difference matrix captures differences in general interactions at protein-ligand interfaces such as van der Waals interactions and pi-pi stacking. For instance, if the atomic pair distance between i,j is 4.0 and 3.0 Å at native and model structure, respectively, the difference value should be -1.0 Å. Distogram prediction (with a binning of 16 from -5 to 5 with the bin size of 1/3) is learned by a cross entropy loss as a auxiliary loss for the expressiveness ^13^.

##### Hydrogen bond difference loss

The hydrogen bond difference matrix is intended to further focus on the hydrogen bonding differences between decoy and experimental structure; SmoothL1 loss is applied as an auxiliary loss.

##### Polar-apolar and Apolar-apolar interaction difference losses

The polar-apolar interaction difference matrix is designed to penalize physically unrealistic interactions often found in docked poses proportional to the polar atom’s partial charge, while the apolar-apolar term encourages hydrophobic interactions; SmoothL1 loss is applied as an auxiliary loss.

##### Information aggregation for the lDDT prediction

The predicted 2-D interaction difference matrices are aggregated into a 1-D vector with the length of the number of ligand atoms. Each value at the vector ranges from zero to one, where a value closer to zero indicates proximity to the near-native structure. Using this interaction embedding along with the aforementioned 1-D ligand embedding vectors (**Figure 3a**), the network finally predicts the complex and atomic lDDT. These values are computed as weighted means of all embedding vectors, where weights are learnable parameters except that for the local atom graph embedding fixed at a constant of 0.1.

#### 3.2.4 Loss functions (complex lDDT, ranking loss)

The final prediction values generated by this network are the complex and atomic lDDT. The original definition of the lDDT is a superposition-free metric assessing local distance disparities among all atoms in a model, with a value 1.0 meaning perfect fit to the answer. We employ this metric to measure the extent to which contacts between the protein and ligand are maintained in decoys compared to their experimental structures. We reasoned this metric is more appropriate than the ligand RMSD (root mean square deviation) popularly used in the area for two reasons: first, native-like contacts between a protein and ligand may be properly captured even at a model-docking context (this doesn’t hold for superposition-based metrics especially when receptor conformation significantly differ from experimental structures); second, it more focuses on overall correctness as opposed to RMSD metric where incorrectness at poorly modeled parts overwhelms the overall value.

Complex lDDT (eq 8 below) appears as a scalar value for each complex, whereas atomic lDDT (eq 9 below) is vector-valued, with its size corresponding to the number of ligand atoms. For the training purpose, thresholds for complex lDDT are set more generously than that for atomic lDDT to distribute labels more evenly:

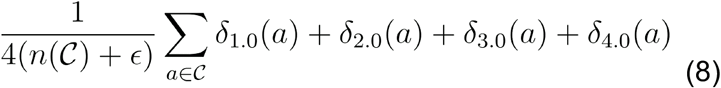

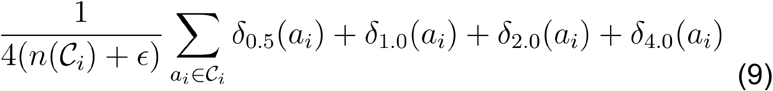

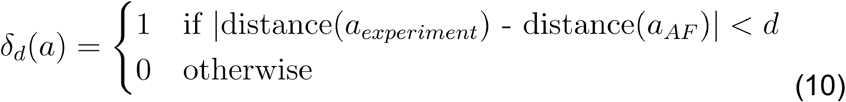

where *C* denotes the number of contacts between the protein and ligand.

In addition to the likeliness loss term, we introduce a ranking loss term. While perfect matching of all values would ideally lead to correct ranking, in reality, accurately predicting all values is challenging, and prediction error can lead to incorrect ranking. Therefore, incorporating a ranking loss term helps address these potential inaccuracies and provides a driving force for improving the ranking performance. During the training step, a batch of graphs from the same complex (i.e. PDBID) and their relative ranking are evaluated altogether. We utilize pairwise ranking loss (implemented as torch.nn.MarginRankingLoss) for all predicted complex lDDT values.

The total loss for the native-likeness prediction network is expressed in Equation (11):

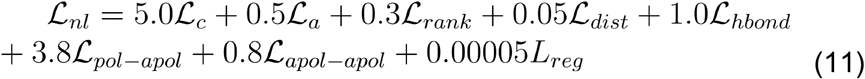

Here *L*_*dist*_, *L*_*hbond*_,*L*_*pol*−*apol*_, and *L*_*apol* − *apol*_ are auxiliary losses aforementioned, while *L*_*c*_ *L*_*a*_, *L*_*rank*_ and *L*_*reg*_ represent the losses associated with complex lDDT, atomic lDDT, ranking, and L2 parameter regularization, respectively. The weights for loss terms are empirically determined without specific target values. Because complex lDDT serves as the major output of the network, higher loss weight is assigned to it compared to that for atomic lDDT. Among interaction terms (hbond, pol-pol, and apo-apol), a higher weight is assigned to the apol-pol term in order to reflect the physically huge energy penalty for buried unsatisfied polar atoms.

#### 3.2.5 Fine tuning strategy for ranking

After the first round training, we performed fine-tuning by increasing the weight for the ranking loss while decreasing weights for rest. The main objective is to increase the network’s selection performance, based on our observation that learning to predict absolute lDDT conflicts with learning ranking especially when the training reaches convergence. The loss weights are adjusted as follows: ranking loss by 10 times, down-weight complex lDDT and atomic lDDT losses by 0.5, and interaction difference matrix losses by 0.1. The effectiveness of fine tuning is reported in the ablation study section of Results.

### 3.3 Binding energy prediction network

#### 3.3.1 Model architecture and Physics-inspired modules

We designed the second sub-network, *the Binding energy prediction network*, which estimates binding energy using explicit force field terms and learned atomic embedding. The network shares the common philosophy with the former network: to apply physical consideration but in a softened form in order to tolerate noises from force field terms. As illustrated in **Figure 3c**, the architecture of the network comprises three primary modules: van der Waals (vdW), electrostatic, and solvation modules. This network solely utilizes an atom graph as input. The network mainly focuses on learning atomic embedding, *h*_*i*_, in which the majority of parameters (in order of tens of thousands) are placed, followed by simple layers containing minimal learnable parameters (in order of hundreds, detailed number of parameters reported in **Figure S6**) at which learned embedding is transformed into energy values through molecular mechanics functional forms (e.g. Coulombic, Generalized Born, etc.). This strategy conceptually corresponds to dynamically assigning atomic parameters (such as vdW radius, partial charge) according to the atomic context, and hence can smartly avoid common artifacts of static atom type assignments typically applied in force fields. Detailed explanations and formal definitions of each module are provided in their respective sections below, and labels for prediction are described in the loss function section.

##### van der Waals module

As represented in the equation below, the van der Waals module comprises two terms: one for attraction energy and the other for repulsion energy. Instead of using the Lennard-Jones term, which returns very stiff repulsion penalty at short distances, each term is approximated using Gaussian functions or higher-order exponential functions on atomic distances governed by a few learnable parameters denoted by the superscript *θ*:

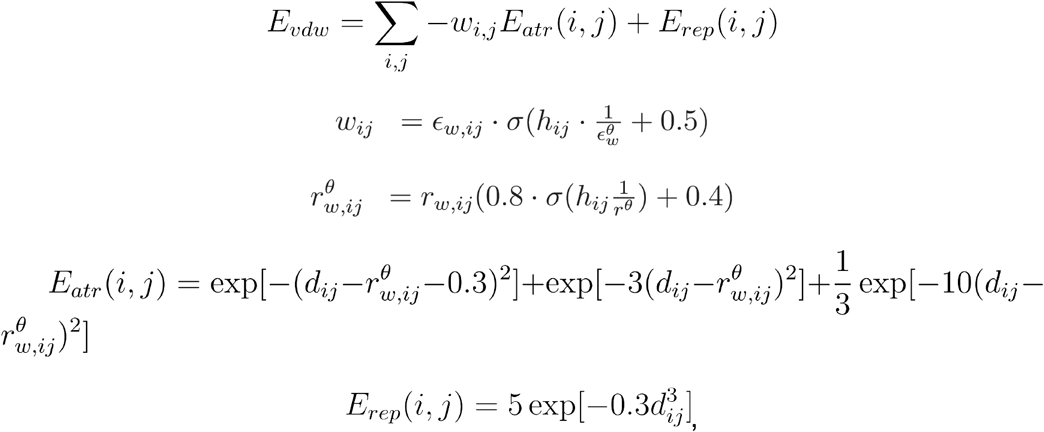

where σ implies sigmoid function, 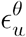 and *r* ^*θ*^ represent learnable parameters that are randomly initialized and are learned to calculate the multiplication factors to the original GenFF ^14^ vdW strength (e_w,ij_) and radius (r_w,ij_), respectively, from the concatenated node embeddings (denoted as *h*_*ij*_). Other constant values are predefined for value-capping or for the curve-fitting.

*Electrostatic module:* The architecture of the electrostatic module is based on the Coulombic potential. Similar to the van der Waals module, this module takes concatenated node embeddings (h_ij_) and incorporates it as an input to estimate context-dependent electrostatic permittivity *(ϵ*^*θ*^) which effectively takes into account environmental effects to electrostatic interactions by a pairwise manner. The electrostatic screening effect is basically described in a distance-dependent manner:

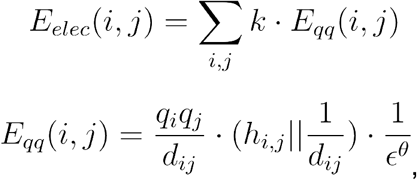

where *k* represents the conversion constant (332.0) for the kcal/mol unit, and *q* denotes the atomic partial charge (taken from Rosetta GenFF ^14^). Note that terms involving multipole moments are not considered here; introducing such expansion may help better capture hydrogen bonding geometry.

##### Solvation module

The solvation module draws inspiration from the Generalized Born (GB) model but simplified to take the self-solvation term alone (note that pair screening term in GB is accounted for by the electrostatic module):

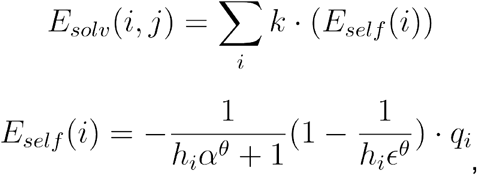

Two learnable parameters, *α* ^*θ*^and *ϵ*^*θ*^, act as scaling factors for learning environment dependent Born radius and electrostatic permittivity, respectively. Other notations remain consistent with those used in the electrostatic module.

#### 3.3.2 Loss functions

The target labels (i.e. per-term physical energy values) for the network are taken from the generalized Rosetta energy function (GenFF) ^14^. Note that approximating physical energy by GenFF is, first because it shows the highest discriminative power for protein-ligand systems among many docking energy functions reported up to date ^14^, and second is for computational efficiency. This approximation can be revisited in the future to use more robust calculations verified for protein-ligand interactions. We calculate the decomposed binding energy in GenFF as the difference between the ligand binding complex and the unbound complex where the ligand is explicitly detached. The total loss for the binding energy prediction network is expressed in the equation below.

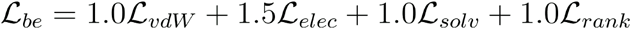

Here, each loss term measures mean-squared-error between the net decomposed energy term value and the predicted value. Additionally, we incorporate ranking loss to address the near-native selection problem. The weights for the loss terms are empirically determined.

### 3.4 Dataset

We utilized the PDBbind v.2020 dataset for the training dataset while excluding the core set for the benchmarking purpose. The training dataset, which should contain model-docked decoys, was generated by running AlphaFold2 (AF2) ^13^. Molecular docking was performed using the preset “flexdock” mode in Rosetta GALigandDock ^14^, or GALD in short hereafter, included in the Rosetta modeling suite. GALD allows for sampling not only ligand conformations but also the side chains of the protein near the bound ligand. Due to the limited performance of AF2 on multimeric proteins, proteins whose the binding site was located at a multimeric interface were excluded for model docking, which comprises about 20% of the entire 19,443 entries, but taken only for generating near-native decoys (see below). 18,094 data points were left after removing targets with unrealistic pockets, which were then split to 17,094 for training and 1,000 for validation. For the training and validation targets, we augmented near-native decoys from two sources: first, a relaxed structure generated by running Rosetta FastRelax ^17^ on the native pose ligand superimposed onto the model; second, self-docked decoys by running docking on the bound-form experimental receptor structures. Total number of decoys were relaxed experimental:self-docked:model-docked = 1:20:80. Details of data curation are provided in the **Supplementary Methods**.

To benchmark the model-docking performance, we employed the PoseBusters benchmark set ^18^. This dataset consists of recently deposited structures, and hence allows us to evaluate the impact from the data redundancy on model performance. The PoseBusters set originally consisted solely of experimental structures. To benchmark DENOISer on the model-docking context, we ran AF2 on this set and sampled docked decoys in the same manner as for training data. We refer to this dataset as the PoseBusters model-docking benchmark dataset. We further experimented with the DENOISer’s performance on broader problems including cross-docking and self-docking problems. 8,302 proteins in the APObind dataset ^19^ and 285 proteins in CASF-2016 (Comparative Assessment of Scoring Functions) dataset ^20^ were taken for the test, respectively. The APObind dataset evaluates docking performance on apo forms, and testing on the set reveals any potential over-fitting issues for the methods that were solely trained on bound poses alone. The original set comprises 13,139 protein pairs, but after excluding those with ligands at inter-chains and also those having ligand preprocessing failures (e.g. RD-kit), final 8,302 complexes were taken. The CASF-2016 dataset, a core set of PDBbind ^21^, was employed to assess self-docking power. These datasets allowed us to evaluate our model’s ability to generalize to cross-docking and self-docking tasks.

## 4. Experiments and Results

Here, we evaluate the performance of DENOISer for model-docking tasks on the PoseBusters benchmark set and cross-docking tasks on the ApoBind set. We chose GALD, AutoDock Vina ^22^, GNINA ^23^, and DiffDock ^24^ for comparison. AutoDock Vina is the most broadly used classic docking tool, while GNINA features an enhanced deep learning scoring function to improve model selection from Vina outputs. DiffDock was selected as the state-of-the-art deep learning method for the given benchmark set. For each method, each PDB ID is subjected to sampling of 100 decoys following the same strategy employed in constructing the training dataset.

### 4.1 Assessment of DENOISer for model-docking

**Table 1** presents the performance of DENOISer on the PoseBusters model-docking benchmark dataset. The left side of the table displays the number of successful PDBs and the success rates for top 1, top 5, and top 10 selections using their respective scoring functions. Success is defined as the selection of a complex structure in the top N (N=1,5,10) with a complex lDDT score exceeding 0.85, which is approximately equivalent to a ligand RMSD of 2.0 Å (**Figure S1**). Success rates are provided in parentheses as two numbers in each cell, representing the success rates for the entire set (left) and for the “valid set” (right) which includes targets only if near-native decoys were sampled.

**Table 1.** PoseBusters model-docking ranking performance. The success is defined as complex lDDT over 0.9 that is approximately equivalent to the ligand RMSD of 2.0 Å. Description at the parentheses of methods represents how receptors are treated: “flexible”, flexible receptor; “rigid”, rigid receptor, “blind”, pocket unknown. *: the success rate for “valid set” in which PDB with no near-native decoys are excluded

| Sampling Method | Original scoring function |  |  | Re-ranked by DENOISer |  |  |
| --- | --- | --- | --- | --- | --- | --- |
|  | Top 1 Complex IDDT | Top 5 Complex IDDT | Top 10 Complex IDDT | Top 1 Complex IDDT | Top 5 Complex IDDT | Top 10 Complex IDDT |
| GALD (flexible) | 97<br>(29.8%, 45.3%*) | 153<br>(46.9%, 71.5%*) | 173<br>(53.1%, 80.8%*) | **124**<br>**(38.0%, 57.9%\*)** | **171**<br>**(52.5%, 79.9%\*)** | **191**<br>**(58.6%, 89.3%\*)** |
| AutoDock Vina (rigid) | 62<br>(19.0%, 35.4%*) | 72<br>(22.1%, 41.1%*) | 105<br>(32.2%, 60.0%*) | 77<br>(23.6%, 44.0%*) | 112<br>(34.4%, 64.0%*) | 128<br>(39.3%, 73.1%*) |
| GNINA (rigid) | 69<br>(21.2%, 49.6%*) | 97<br>(29.8%, 69.8%*) | 114<br>(35.0%, 82.0%*) | 61<br>(18.7%, 43.9%*) | 84<br>(25.8%, 60.4%*) | 101<br>(31.0%, 72.7%*) |
| DiffDock (blind, rigid) | **127**<br>**(39.0%, 63.0%\*)** | **156**<br>**(47.9%, 77.2%\*)** | 169<br>(51.8%, 83.7%*) | 124<br>(38.0%, 61.4%*) | 153<br>(46.9%, 75.7%*) | 167<br>(51.2%, 82.7%*) |

The best performance is achieved by DENOISer when combined with sampled decoys using GALD. The side-chain sampling in GALD is likely to benefit generating a high-quality pool for selecting near-native protein-ligand complexes in the model-docking scenario. Through the implementation of the re-ranking strategy, we observe a notable increase in the success rate over GALD by 14.6% (from 45.3% to 57.9%) when only valid targets are counted (increment of 8.2% for the entire set). Although this increment may not be dramatic, it underscores the potential utility of the re-ranking strategy in model docking.

We further analyze the results based on the receptor redundancy to the training data (PDBbind 2020) and the physical plausibility of docked poses (**Figure 4)**. The three bars represent success rates broken down by maximum sequence identity to any receptor in the training dataset. Consistent with the PoseBusters set paper ^18^, Diffdock, a deep learning method, exhibits a noticeable decrease in success rates with the low sequence identity to the training data (note that DENOISer and DiffDock share the common training set of PDBbind 2020). However, DENOISer, as well as the original GALD, show relatively robust success rates across different levels of sequence identity. Moreover, the GALD outputs are generally much more physically valid (darkest bars in the figure; < 1% failure) compared to those generated by DiffDock (>60% failure). Overall, while DiffDock shows similar performance on well-known receptors, DENOISer outperforms it in terms of physical validity as well as transferability to novel targets.

**Figure 4.**
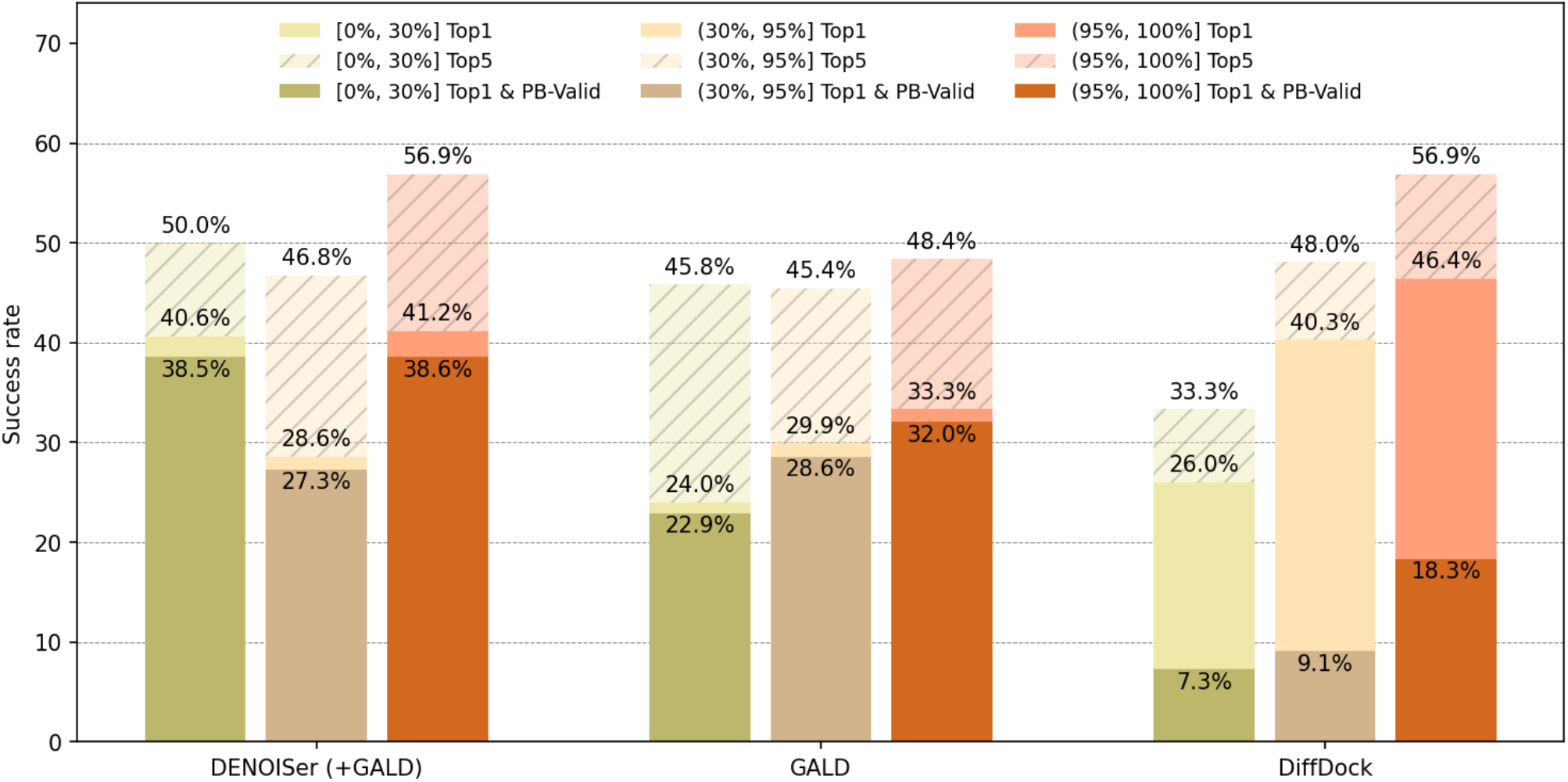
Performance on PoseBusters set stratified by sequence identity relative to the PDBbind general set v2020. The striped bar indicates the performance of the top 5 selections; light bar and dark bar indicates top-1 success and [top-1 success & physically valid], respectively. Success is defined as complex lDDT over 0.85. Physical validness was calculated using PoseBusters tools.

To understand the origin of success, we conducted detailed case studies. **Figure 5** illustrates examples of near-native solutions identified through DENOISer (a,c) but failed by GALD score (b,d). It is evident that several differences exist between the experimental and predicted structures, which can significantly impact model-docking performance. Looking at the panel a-b, we speculate that the highlighted structural discrepancy may displace the ligand from the binding site and cause decreased vdW packing; however, DENOISer is more tolerable to such structural variations and can favor overall solvation effect to accurately select the near-native solution (originally ranked as 46th). **Figure 5c**,**d** shows another case where DENOISer could make a correct selection; subtle structural deviations at the structurally dynamic region of the protein, frequently found in AF2 models (highlighted by darker colors), cause the decision difference. Overall, while the GALD scoring function (GenFF, and presumably the majority of docking score functions) focus on perfect vdW packing even at structurally variable regions, DENOISer rather highly penalizes the unfilled void or solvation effects. We argue that this discrepancy cannot be simply resolved within docking score functions by re-weighting terms; a more holistic view learned via deep learning should have enabled this complicated balancing of terms within model docking contexts.

**Figure 5.**
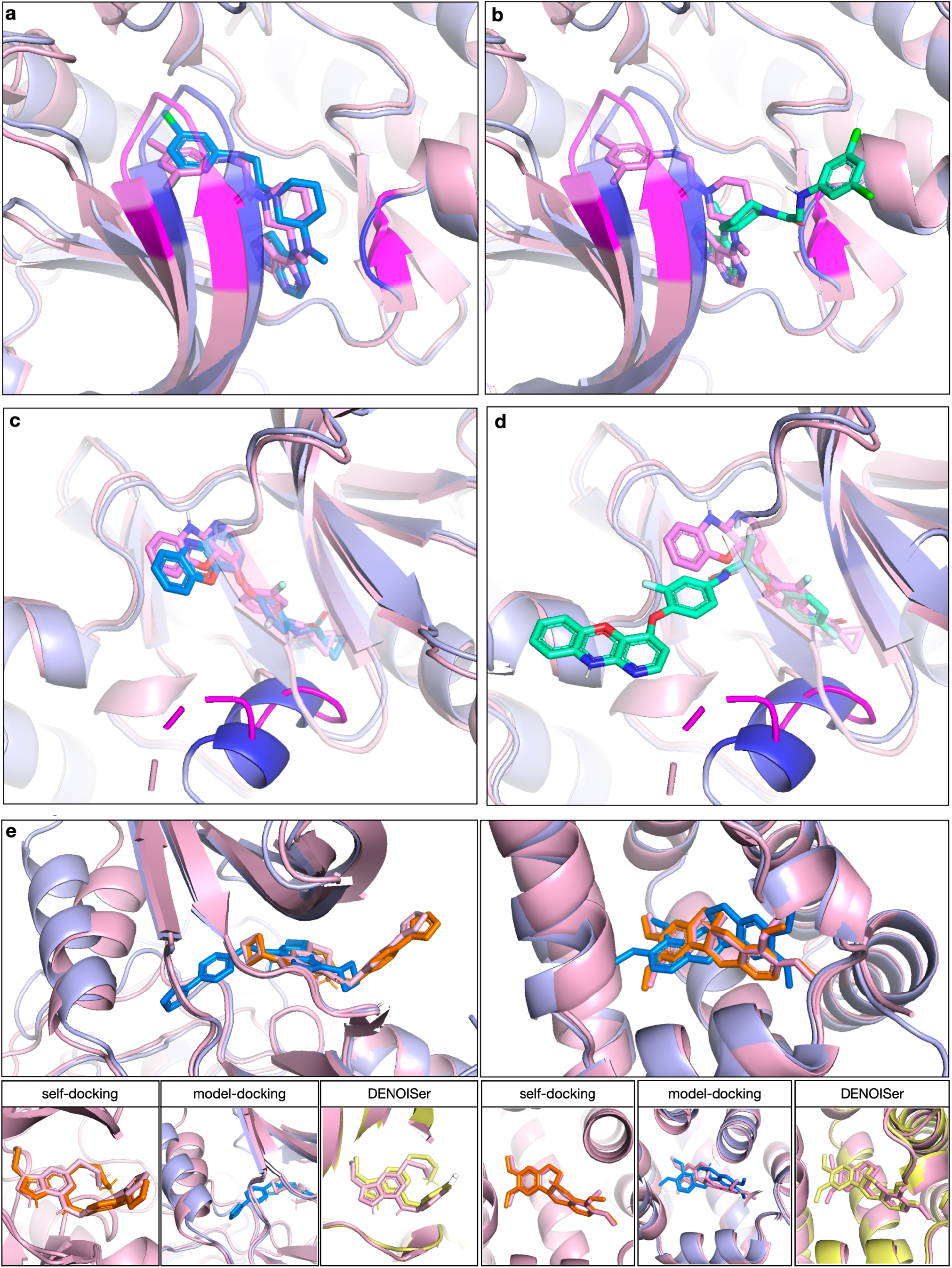
Examples of near-native solutions selected by DENOISer. (a)∼(d) pink: X-ray crystallography, blue: AF2 predicted and re-ranked by DENOISer; cyan: AF2 predicted and selected by GALD. Major structural errors near the ligand are highlighted in darker colors. (a) PDB (8FLV) 1st selected by DENOISer. (b) 1st selected by docking score of GALD. (c) PDB (7V3S) 1st selected by DENOISer. (d) 1st selected by docking score of GALD. (e) Cases where self-docking succeeded but model-docking failed with GALD. The panels below show the first selected structures when performing self-docking (left) and model-docking with GALD (middle) and the selection by DENOISer (right). blue: AF2 predicted, orange: self-docking.

Finally, to see how crucial the receptor structure impacts docking results, we compared self-docking, model-docking by GALD, and re-ranked by DENOISer in **Figure 5e**. These examples highlight how commonly typical docking tools should have failed to recover native poses in model docking which originally succeeded in self-docking, highlighting the usefulness of DENOISer in processing model-docking outputs.

### 4.2 The network is generalizable to different docking data and to broader docking tasks

In **Table 1**, the dependency of DENOISer on sampling tools is also compared. The success rates for the valid set (target containing at least one near-native) do not exhibit significant dependencies on the docking tools used for sampling, despite the fact that DENOISer was trained solely on GALD generated data. This demonstrates DENOISer doesn’t have a bias toward GALD decoys and can be combined with other sampling methods. The top 10 performance on GALD decoys is still higher than the rest, suggesting broader space covered by GALD. Interestingly, GNINA performed worst among tested models, presumably because training a deep learning model solely on self-docking examples is not generalizable enough to model-docked structures. Re-scoing DiffDock also didn’t help much because the outputs were already too similar to each other.

Cross-docking presents another formidable challenge in molecular docking research, as highlighted in numerous studies ^25–28^. As an example, the top-1 success rate of SMINA was 57.9% in self-docking, but dropped to 13.7% in apo-form cross-docking. To evaluate the applicability of our model to cross-docking tasks, we employed the APObind benchmark set consisting of 8,302 complexes and sampled decoys using GALD. We then compared our model’s performance with GALD’s original docking scoring and to other tools.

**Table 2** shows that DENOISer significantly enhances the success rate of GALD on cross-docking, achieving up to an 21.5% increase (from 36.5% to 58.0%) for the top 1 selection and a 21.1% increment for top 5 selection when only valid targets are considered. Furthermore, results on the core set of APObind, which exhibits less protein sequence redundancy compared to the general and refined sets of PDBbind, show a similar trend. DENOISer also showed reasonable performance on self-docking cases; tested on CASF-2016, DENOISer was good at ranking in the top-3, but less accurate for selecting top-1. Overall, these findings suggest that our re-ranking strategy can be effectively applied to the challenges of cross-docking, and will be more effective when near-natives can be sampled, highlighting the potential of our model in this task.

**Table 2.** Expansion to other tasks: cross-docking and self-docking. DENOISer re-ranked the structures sampled from GALD. Above are the benchmark results for the APObind set for cross-docking task evaluation, and below are the results for the CASF-2016 set for self-docking task evaluation. Because the CASF-2016 set has a near-native structure in all decoys, success rates are not indicated. *: the success rate excluding PDB which has no near-native decoys 1) The relatively small total number is due to RDKit’s failure to read the file.

| Cross-docking (APObind) |  |  |  |  |
| --- | --- | --- | --- | --- |
| Method | Top 1 | Top 5 | Top 10 | Run without processing error |
| GALD | 1,535<br>(18.5%, 36.5%*) | 2,477<br>(29.9%, 58.9%*) | 2,959<br>(35.7%, 70.4%*) | 8,294/8,302 |
| Re-ranked by DENOISer (+GALD) | **2,435**<br>**(29.4%, 58.0%\*)** | **3,361**<br>**(40.5%, 80.0%\*)** | **3,676**<br>**(44.3%, 87.5%\*)** | 8,294/8,302 |
| AutoDock Vina | 1,189<br>(14.7%, 33.4%) | 1,569<br>(19.4%, 44.1%) | 2,074<br>(25.7%, 58.3%) | 8,067/8,302 |
| GNINA | 1,286<br>(17.0%, 49.1%*) | 1,823<br>(24.1%, 69.6%*) | 2,121<br>(28.1%, 81.0%*) | 7,559/8,302 |
| DiffDock† | 2,363<br>(40.1%, 71.9%) | 2,761<br>(46.9%, 84.0%) | 2,346<br>(49.4%, 88.5%) | 5,888/6,481<sup>1)</sup> |
| Core set |  |  |  |  |
| GALD | 37<br>(21.9%, 38.1%*) | 63<br>(37.3%, 64.9%*) | 71<br>(42.0%, 73.2%*) | 169/169 |
| Re-ranked by DENOISer (+GALD) | **51**<br>**(30.2%, 52.6%\*)** | **74**<br>**(43.8%, 76.3%\*)** | **86**<br>**(50.9%, 88.7%\*)** | 169/169 |
| AutoDock Vina | 15<br>(9.0%, 19.7%) | 25<br>(15.1%, 32.9%) | 40<br>(24.1%, 52.6%) | 166/169 |
| GNINA | 33<br>(22.6%, 51.6%*) | 48<br>(32.9%, 75.0%*) | 52<br>(35.6%, 81.3%*) | 146/169 |
| DiffDock† | 44<br>(35.2%, 71.0%*) | 50<br>(40.0%, 80.6%*) | 52<br>(41.6%, 83.9%*) | 125/129<sup>1)</sup> |

|  | Top 1 Ligand RMSD | Top 2 Ligand RMSD | Top 3 Ligand RMSD |
| --- | --- | --- | --- |
| Re-ranked by<br>DENOISer (Ligand<br>RMSD under 2.0) | 217<br>(76.14%) | 237<br>(83.2%) | 280<br>(98.2%) |

### 4.3 Confident docked poses can be selected using predicted values

If an accuracy estimator can provide a confidence measure alongside the prediction, this measure can guide users in making more informed decisions about whether to trust the docking pose. By grouping prediction results, we see improved selection quality for those targets surpassing the predicted native-likeness of 0.75 (**Fig 6a**): chance to pick a near-native pose as top1 increases to 60%, compared to 38.0% when no such condition was applied. This observation highlights the advantage of our network over typical docking scores, where a clear threshold is lacking due to the weak correlation between pose accuracy and score values.

**Figure 6.**
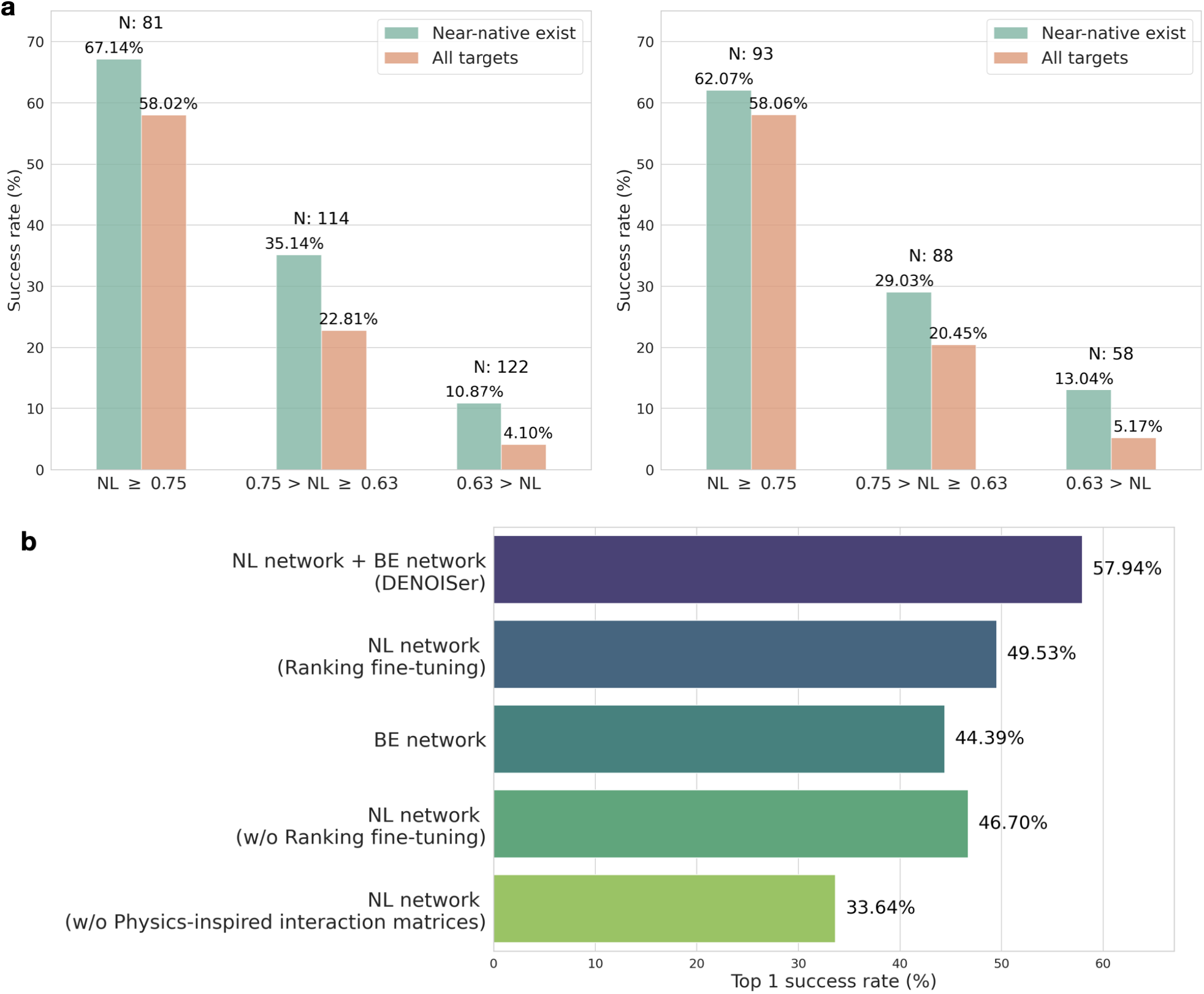
Effect of DENOISer confidence metric on the success rate and ablation study on PoseBusters model-docking dataset. (a) Top-1 success rate by GALD docking score with DENOISer confidence metric applied across different criteria. Left: PoseBusters with generated ligands. Right: PDBbind test set (model docking). Statistics are shown for a subset where near-native solutions existed in the pool (green bars) or all dataset regardless of it (orange bars). (b) Network ablation study on PoseBusters model-docking dataset. NL network: Native-likeness prediction network, BE network: Binding energy prediction network.

### 4.4 Ablation study highlights importance of the physics-inspired architecture and the model consensus ranking

We conducted an ablation study to confirm the effect of the individual contributions of components within DENOISer. **Figure 6b** depicts the success rates achieved using the PoseBusters model docking set. The top bar is the result with our model, which uses both the Native-likeness (NL) prediction network and the Binding energy (BE) prediction network. The second and third bars are the stand-alone performances of the Native-likeness prediction network and Binding energy prediction network, respectively. Although the performance of the Native-likeness prediction network is relatively better, we can see that the consensus score using the two sub-networks is complementary and better. Additionally, fine-tuning of the Native-likeness prediction network brought further enhancement. Notably, the absence of a physical interaction difference matrix results in the poorest performance, underscoring the importance of physical inductive biases encoded in the network architecture.

## Discussion

To check how robust DENOISer performs upon receptor structure variation, we generated interpolated receptor structures for the PoseBusters set between experimental and AF2 model structures, with subsequent docking onto those. If DENOISer was to memorize model and experimental receptor structures, but not else, the performance will peak at both points; in contrast if the model is broadly applicable to various receptor conformations, its performance should gradually decay. **Figure S3** shows the robustness of our model against structural errors in the predicted proteins for these datasets – while GALD drops from 83.4% to 62.0%, DENOISer’s performance stays quite steady from 86.5% to 75.4% with a gradual decay. This observation suggests that our model exhibits a capacity to select near-native solutions at a reasonable range of error levels.

One of the caveats of our network is its dependence on high quality samples. If no near-native sample exists, then the network may eventually fail. Checking the quality of ligand samples first, we repeated our experiments by generating ligands from scratch using CORINA v5.0 (instead of borrowing from experimental structures). We observed difference in the discrimination performance especially for the PoseBusters set, while such discrepancy was negligible for the PDBbind test set (**Table S4**). Investigating ligand structures, more failures were found from the PoseBusters set originating from the ring geometry, wrong tautomer assignment, or cis/trans flip, suggesting difference in the characteristics between the sets and also need for improved ligand conformation (or tautomer) generation. Second, to see the network’s ability upon complex structure quality, we relaxed experimental structures in the PoseBusters set, adding near-native structures for 65 more targets (total 279 of 326). The rest 47 targets remained unsampled due to substantial protein backbone differences at AlphaFold2 models or other missing cofactors/metals. The same issue was present for a large portion of new 65 valid targets (**Figure S4**), only 43 of which were selected as top-1 (**Table S2**). Although the network was trained without such molecules so as to implicitly learn neighboring molecules, those types of problems remained relatively more challenging. However, there were other cases DENOISer clearly failed, including those having explicit water bridges (**Supplementary Methods**), demonstrating need for further method development. We anticipate that further improvement of the binding energy prediction network, combined with improved ligand conformation sampling, would improve the model’s ability to address these complexities. Overall, while the success rate of DENOISer increases with the addition of near-native data, there remains a gap to be addressed.

In this work, we presented DENOISer, a deep evaluation network specialized for selecting correct docking solutions in model-docked scenarios. It was demonstrated that physical consideration while designing the network brought key contributions to its improved performance over existing docking score functions. Meanwhile, its limited performance originating from the sampling bottleneck clearly suggests the necessity of joint sampling of receptor and ligands, which is recently becoming an active research direction ^29–32^. With the progress in such improved sampling, we anticipate a corresponding increase in the utility of DENOISer.

## Supporting information

The Supporting Information is available free of charge via the Internet at http://pubs.acs.org/.

## Supplementary Information

### Supplementary Tables

**Table S1.**
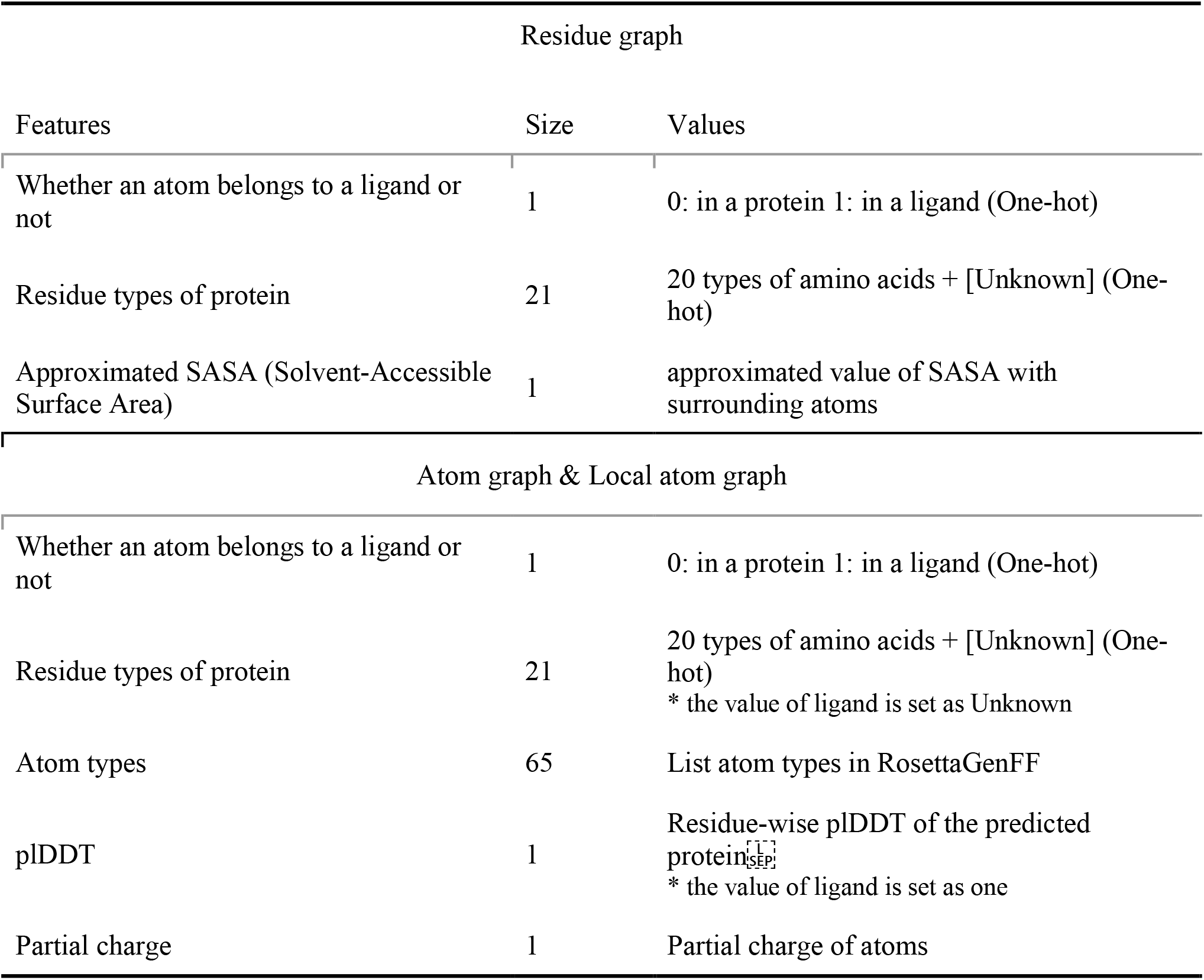
Initial node and edge features.

**Table S2.** Effect of adding near-native solutions to our model selections. Our model re-ranked the structures sampled from GALD. The success is defined as Complex lDDT over 0.85

| Method | Top 1 Complex IDDT | Top 5 Complex IDDT | Top 10 Complex IDDT |
| --- | --- | --- | --- |
| Before adding near-native structures, re-ranked by DENOISer (+GALD) | 124<br>(38.0%, 57.9%*) | 171<br>(52.5%, 79.9%*) | 191<br>(58.6%, 89.3%*) |
| After adding near-native structures, re-ranked by DENOISer (+GALD) | 167<br>(51.2%, 59.9%*) | 219<br>(67.2%, 78.5%*) | 242<br>(74.2%, 86.7%*) |

\*: the success rate excluding PDB which has no near-native decoys

**Table S3.** Hyperparameters.

| Native-likeness prediction network |  |
| --- | --- |
| Parameters | Values |
| Learning rate | $6 \times 10^{-4}$ |
| Number of layers in EGNN block of Graph Embedding Module | Residue graph: 4<br>Atom graph: 6<br>Local atom graph: 3 |
| Graph edge connection | Based on Euclidean distance<br>Residue graph: 8.0Å<br>Atom graph: 4.5Å<br>Local atom graph: 3.0Å |
| Dropout rate | 0.3 |
| Projection dimension of Projection block | 20 |
| Hidden dimension of EGNN block | 32 |
| Dimension of node embeddings as output of Graph Embedding Module | 32 |
| Hidden dimension of physics-inspired interaction difference matrices | 16 |
| Optimization algorithm | AdamW |
| Binding energy prediction network |  |
| Parameters | Values |
| Learning rate | $5 \times 10^{-5}$ |
| Number of layers in EGNN block | Atom graph: 6 |
| Graph edge connection | Based on Euclidean distance<br>Atom graph: 4.5Å |
| Dropout rate | 0.3 |
| Projection dimension of Projection block | 20 |
| Hidden dimension of EGNN block | 32 |
| Dimension of node embeddings | 32 |
| Number of parameters in van der Waals module | $\epsilon_w^\theta$: 64<br>$r^\theta$: 64 |
| Number of parameters in Electrostatic module | $\epsilon^\theta$: 64 |
| Number of parameters in Solvation module | $\alpha^\theta$: 32<br>$\epsilon^\theta$: 32 |
| Optimization algorithm | AdamW |

**Table S4.**
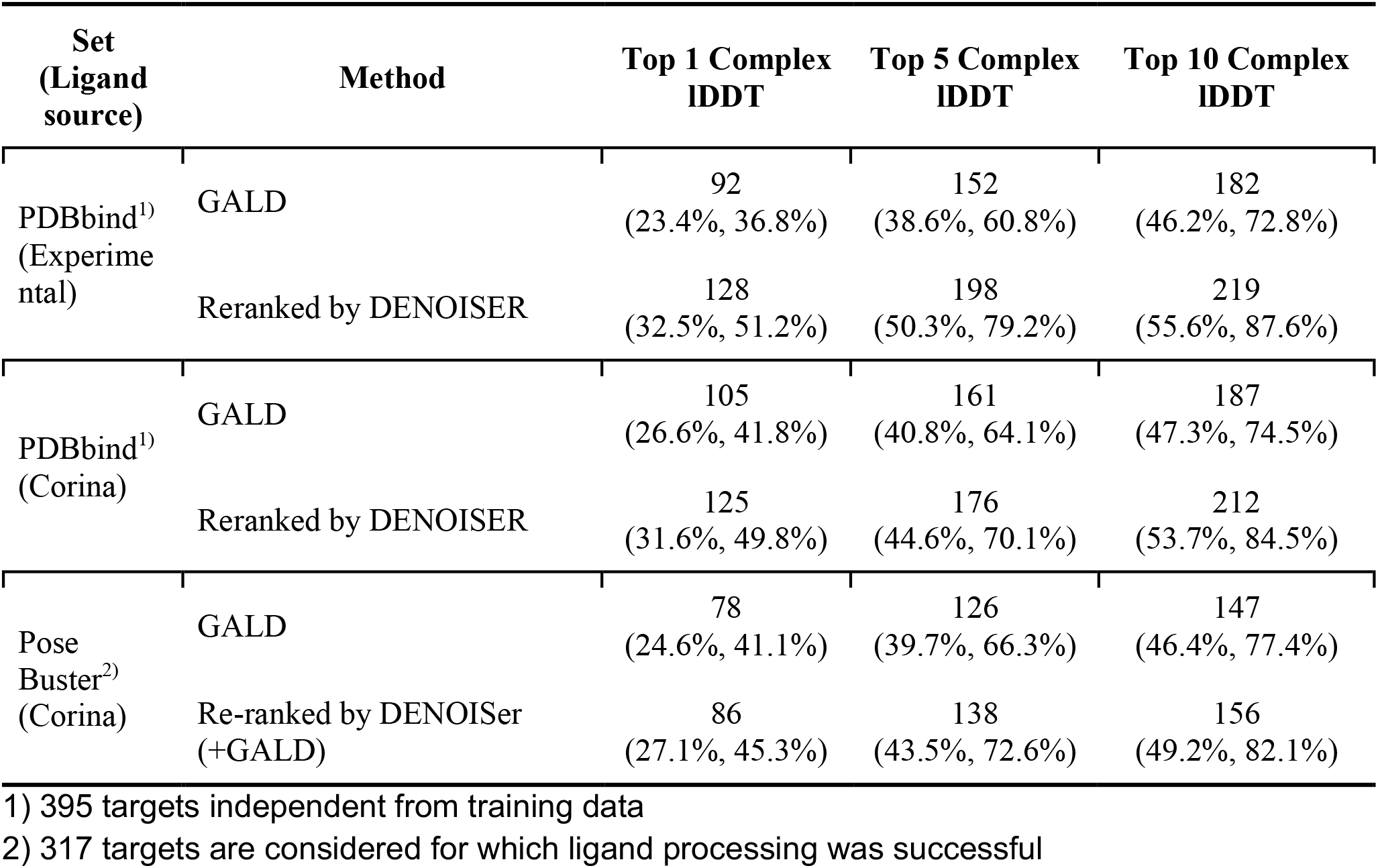
Model-docking ranking performance with generated ligands (Complex lDDT ≥ 0.85).

**Table S5.** PB-valid for Top 1 success cases on APObind benchmark dataset.

|  | GALD | DENOISer<br>(+GALD) | AutoDock<br>Vina | GNINA | DiffDock |
| --- | --- | --- | --- | --- | --- |
| Percentage of<br>PB-valid | 98.9% | 98.5% | 94.9% | 90.8% | 40.0% |

### Supplementary Figures

**Figure S1.**
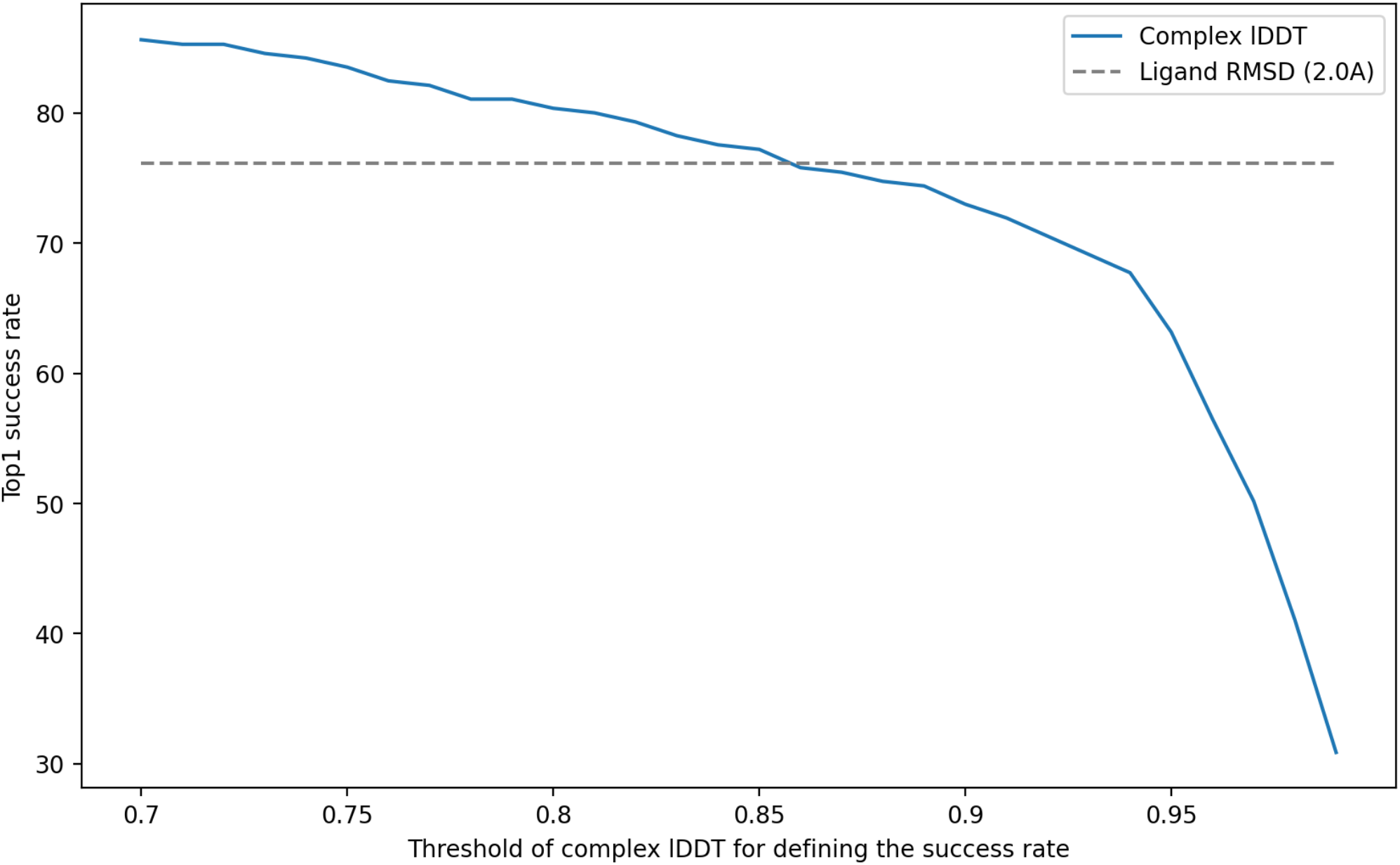
The equivalent relationship between Ligand RMSD and Complex lDDT. The blue curve illustrates the success rate based on complex lDDT with given thresholds (x-axis) in the CASF-2016 dataset. The dashed gray line represents the success rate with Ligand RMSD cutoff of 2.0 Å. Although the intersection point occurs slightly above 0.85, we set a slightly stricter condition, defining success as ‘over 0.85 complex lDDT’.

**Figure S2.**
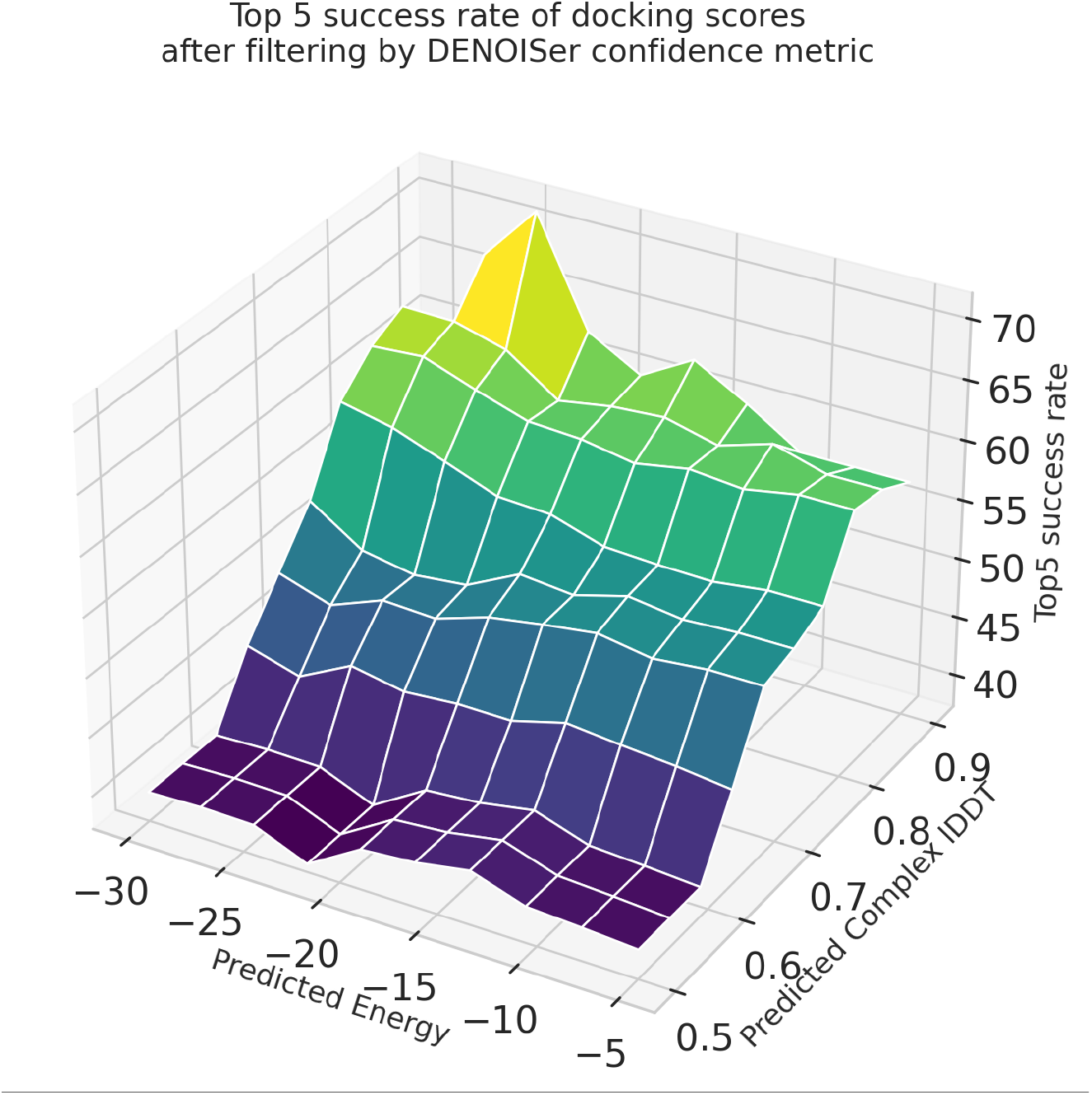
Confidence estimation from the ranking consistency from two sub-networks.

**Figure S3.**
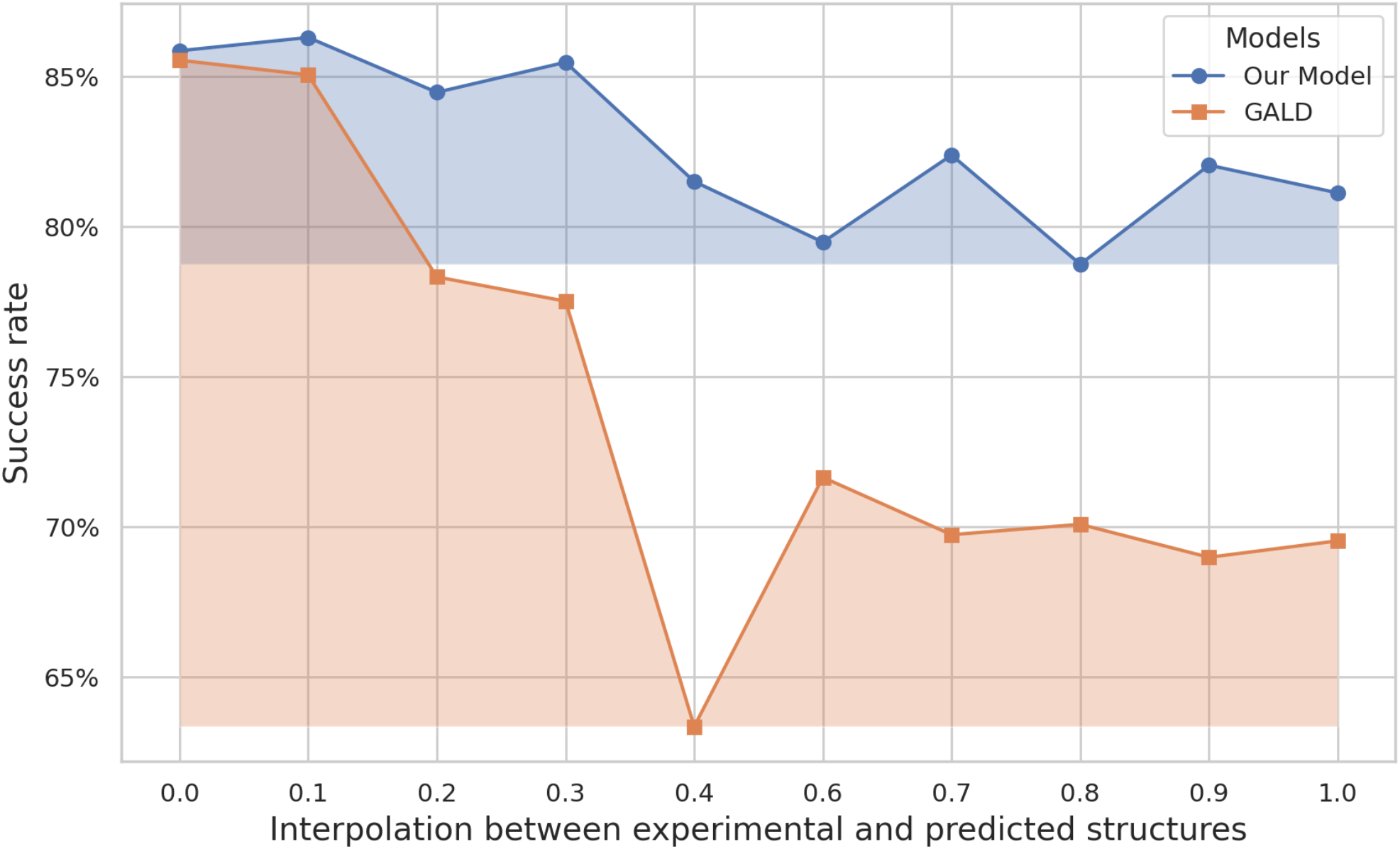
Robustness of our model to protein structural error. The figure depicts the success rate of docking on the interpolated protein structures, which lie between the experimental and predicted structures. The x-axis represents the parameter λ in the equation λ * predicted + (1-λ) * experimental.

**Figure S4.**
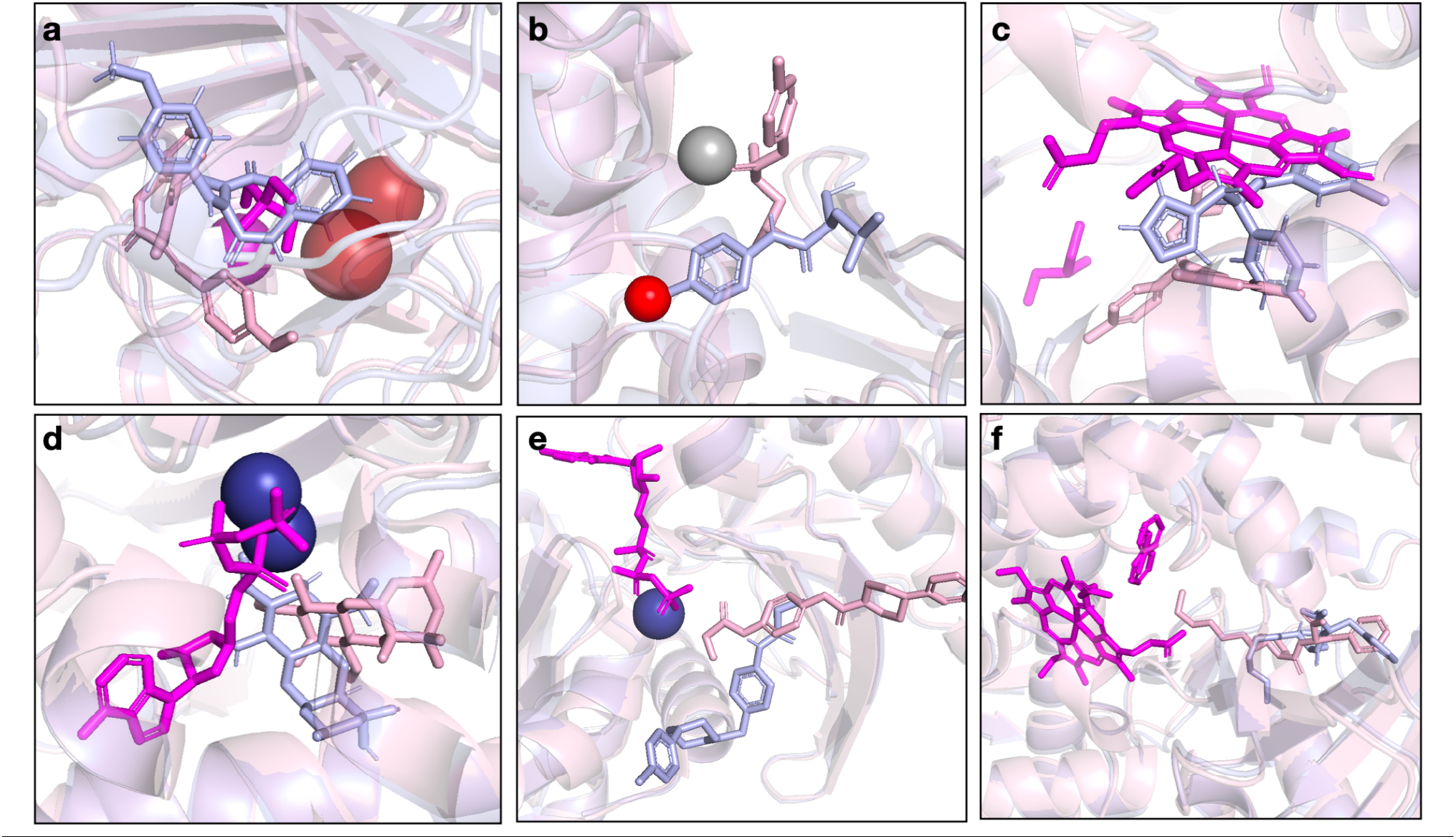
Examples of structures that cannot be selected by our model when a near-native structure is added. pink: X-ray crystallography, blue: AF2 predicted, magenta: cofactors, red: water, gray, cobalt: metal

**Figure S5.**
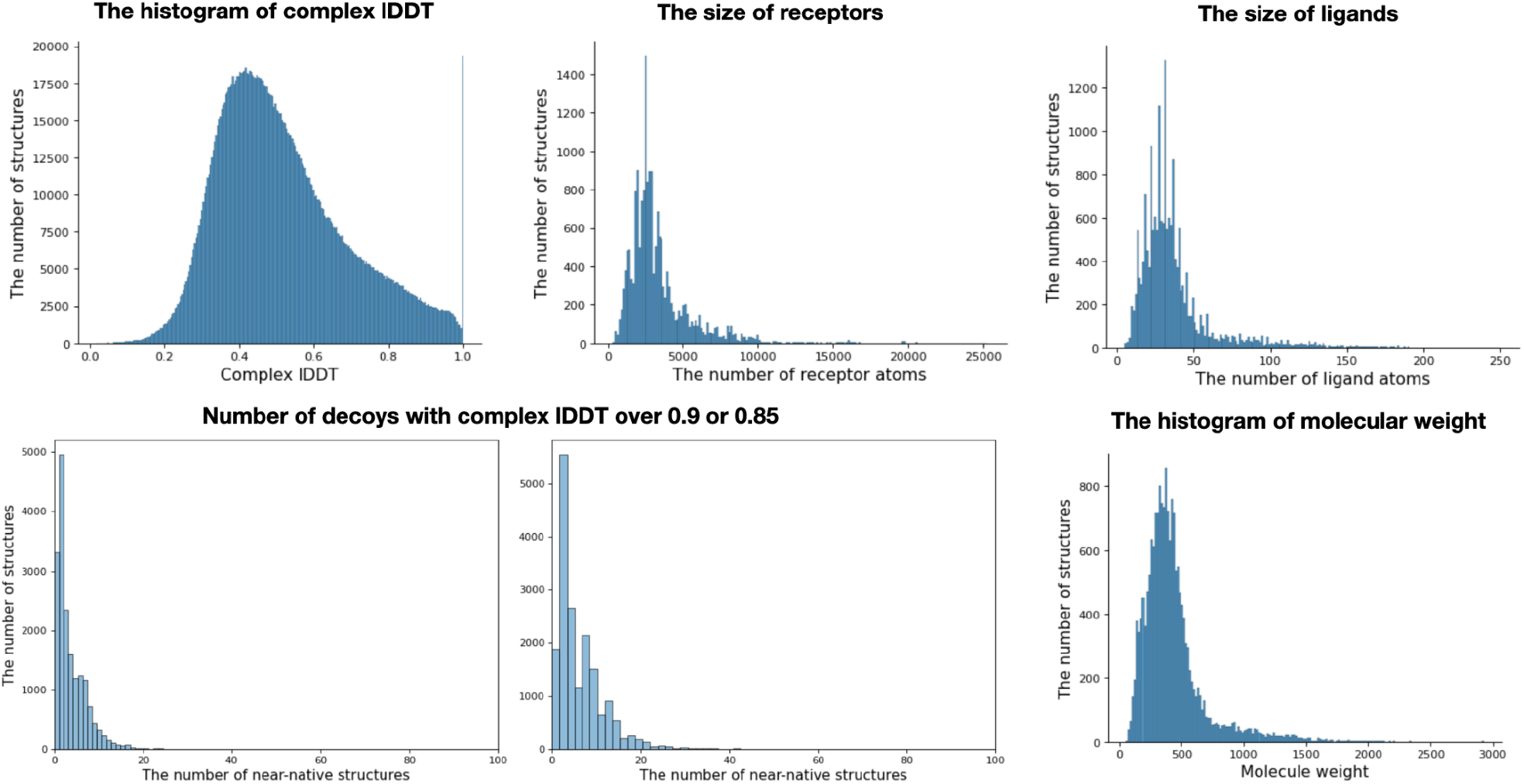
Descriptive statistics of the training data.

**Figure S6.**
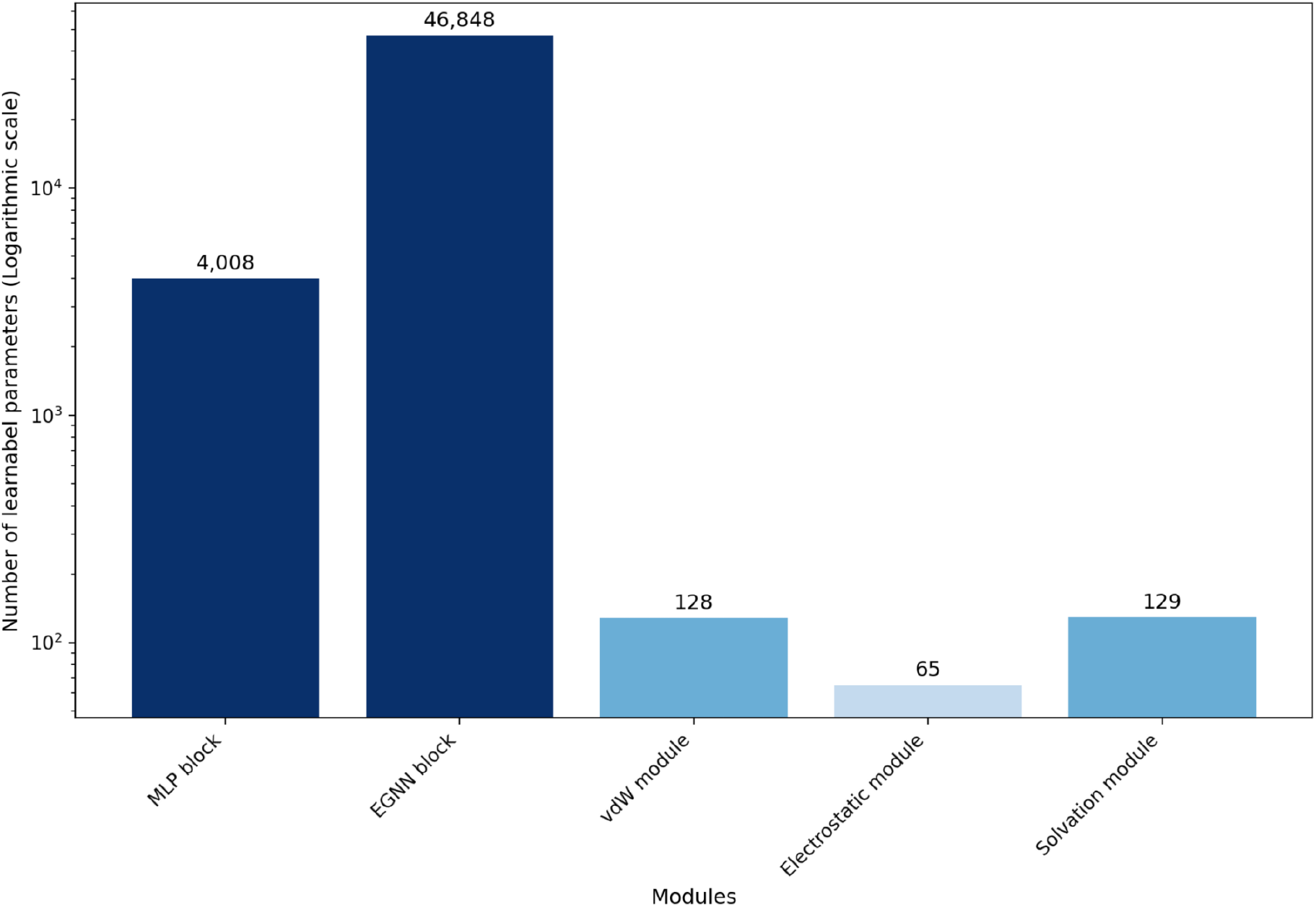
The number of learnable parameters in Binding energy prediction network. The blue color intensity highlights the magnitude of the values. The y-axis is scaled logarithmically. Most of the parameters in the network are contained in EGNN blocks (91.54% of the total parameters).

### Supplementary Method

#### A Training details

##### A.1 Data curation and decoy sampling

The PDBbind v.2020 dataset is used as the training dataset and we run AlphaFold2 exclusively to the complexes that have a binding site localized to a single chain. Complexes with ligands located at interchain binding sites or interacting with multiple protein chains were excluded due to the lower performance of AlphaFold2 Multimer, which struggles to sample near-native structures when conducting model docking with these predicted multi-chain proteins. For instance, it is common for the positions of each chain to be misaligned, resulting in an open binding site where the ligand is supposed to interact. This misalignment can introduce noise into the training set, thereby disrupting the training process and leading to overfitting.

However, simply excluding the data could be limited, due to the data scarcity problem, which is common in this domain. Therefore, we use crystal structures and conduct self-docking for these complexes instead of using predicted structures and model-docking. This strategy can induce intrinsic data bias due to differences in data generation methods. We conduct self-docking to the remaining data and we adjust the ratio of model docking and self-docking as 80: 20. Additionally, near-native structures were added to each PDB id as described in 3.4.

We excluded NMR structures and X-ray crystallography structures with resolutions higher than 3.0 Å. To minimize noise, we set TM-score cutoffs of 0.8 for the overall protein structure and 0.75 for the binding site between the native and predicted protein structures. Trivial ligands, such as benzene, oxalate, dichloroethane, and malonic acid, were also excluded.

At the training stage, all the clusters are sampled once per epoch and the target is sampled randomly from each cluster but with different combinations of decoys. The cluster is constructed using CD-HIT ^33^ with a 70% protein sequence identity. 4 decoys from the same complex are randomly sampled from the pool and sent to the model altogether as a minibatch.

##### A.2 Input features

As illustrated in Supplementary Table 3, the initial node and edge features entered into the model were set. In the residue and atom graph, the protein and ligand nodes are incorporated together into a complex 3D graph, the feature to distinguish them is added. The residue type is defined as input feature, whereas the ligand is classified as unknown. The atom type in the atom graph follows the Rosetta GenFF, allowing for the consideration of various types beyond the simple use of atom elements. Edge features include distance information to explicitly utilize geometric features. Further details are provided in the table.

##### A.3 Residue and atom graph construction

The graph representation consists of both residue-level and atom-level graphs, with the information on each node and edge connection detailed in section 3.2.2. The intended roles of these graphs differ according to their resolution. The atom-level graph includes only the binding site near the ligand due to the memory issue, enabling the model to learn interactions between atoms due to its high resolution. However, the limited information from this localized graph can constrain the model’s learning and potentially lead to overfitting. Furthermore, protein-ligand interactions involve not only short-range but also long-range interactions, which are omitted in high-resolution graphs due to the truncated protein-ligand complex structure.

To address this, a coarse resolution residue graph is introduced, allowing the model to learn long-range interactions and consider the overall protein structure. This multi-resolution approach helps the model to capture both local atomic interactions and global structural features, enhancing its learning capability.

##### A.4 Weight sampling for training and stop gradient

During model training, we considered the distribution of the dataset. Initially, proteins were clustered based on a sequence identity threshold of 70%. These clusters are then included in the training process with uniform probability, ensuring that entries within the same cluster were selected with equal probability. Consequently, if a cluster contained many PDB IDs, each entry had a lower probability of being selected in the training process.

Near-native structures in training dataset are relatively rare because they are difficult to sample. Uniformly sampling all decoys for training can result in being fewer selected of near-native structures, creating an imbalanced training dataset and hindering model performance. To address this, we applied up sampling based on the complex lDDT, giving preference to near-native structures and alleviating the label imbalance issue in the dataset.

When training the model, we used physics-inspired interaction difference matrices to create interaction embedding vectors for lDDT prediction. The goal is to accurately align the loss functions defined in the equations, ensuring that interactions between native structures and decoys are correctly decomposed. Ideally, backpropagation from lDDT to the physics-inspired interaction difference matrices need to be avoided. Thus, we applied a stop-gradient to the interaction embedding vector before using it for lDDT prediction, preventing gradients from flowing back into the interaction difference matrices.

##### A.5 Energy calculator with the binding energy prediction network

The binding energy computed with Rosetta is used as the label for the binding energy prediction network. For simplification, the energy calculation is performed after removing the ligand from the original protein-ligand complex structure, with the difference of the energy being defined as the binding energy. This approach may result in an overestimation of the more precise binding energy. Nevertheless, due to the use of consensus scores from two models in DENOISer, we selected it for its simplicity and efficiency. For the labeling, the van der Waals energy is calculated as the sum of two terms from GenFF: {fa_atr and fa_rep}; the electrostatic energy as the sum of three terms: {fa_elec, hbond_sc, hbond_bb_sc}; and the solvation energy as the sum of {fa_sol, lk_ball, lk_ball_iso, lk_ball_bridge, and lk_ball_bridge_uncpl}.

To explain each term, fa_atr, fa_rep denote Lennard-Jones attractive and repulsion energy, respectively. fa_sol denotes Lazaridis-Karplus solvation energy ^34^, which is the energy of a molecule in implicit solvent using a Gaussian-exclusion solvent model. lk_ball, lk_ball_iso, lk_ball_bridge, and lk_ball_bridge_uncpl are the terms for solvation effect. Anisotropic solvation effect is considered using the previous two terms, and the effect of bridging waters is indirectly considered through the lk_ball_bridge term. However, as this is an approximate method, its limitations are discussed in the D. Limitation section.

#### B Model details

##### B.1 Physics-inspired interaction difference matrices in Native-likeness prediction network

###### Distance difference matrix

The distance difference matrix describes the distogram of clamped distances between the protein and ligand. Equation (1) represents the loss term associated with this matrix, where *y*_*ij*_ denotes the ground truth matrix of the distogram in the experimental structure, *p*_*ij*_ denotes the predicted matrix generated by our model, and *d*_*ij*_ denotes the Euclidean distance between the *i* th atom of the ligand and the *j*th atom of the protein, as indicated in Equation (2). To constrain the difference value of the distance, we limit it to an absolute value of 5. This is achieved by computing *y* using the equation *y*_*ij*_, = *bin (clamp(a*_*ij*_)), where *clamp(x) = max (*5,*min(x −* 5). The bin size is set to 16.

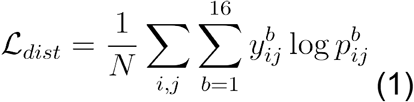

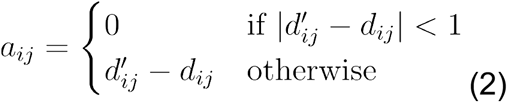

###### Hydrogen bond difference matrix

The hydrogen bond difference matrix elucidates the difference in hydrogen bonding patterns between decoy and experimental structures. Analogous to the distance difference matrix, this matrix serves as a set of predicted values. Notably, the difference is defined with a function *f(d)* dependent solely on the hydrogen bonding distance. The intuition behind not using more complex functions to describe hydrogen bonding is as follows: the primary objective of this network is not to predict precise binding energies. Instead, predicting binding energy is the role of the binding energy prediction network described below. The information contained within this matrix helps improve the network’s understanding of the specific hydrogen bonding characteristics typical of near-native structures. Therefore, we opt for a simplified function to give the network greater flexibility. To compute the loss, we employ a smooth L1 loss function while masking other interactions. *L*_*i*_ and *P*_*j*_ denote an atom of ligand or protein, respectively.

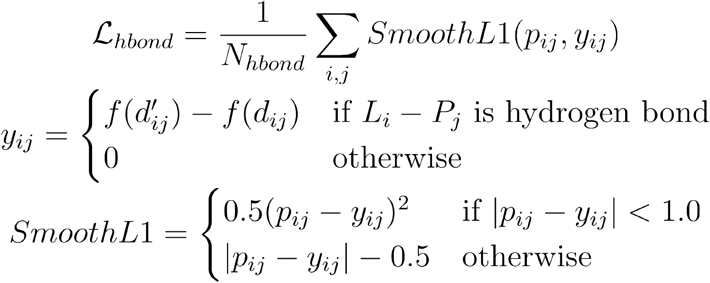

where f is a scaling function, which is *max* (1,*min* (1.8*/x + ϵ)*^4^, 0) and *ϵ* is a small number for computational stability, depending on the distance between atoms.

###### Polar-apolar and Apolar-apolar interaction difference matrix

The polar-apolar and apolar-apolar interaction difference matrices are defined by nearly identical functions. The key distinction lies in the polar-apolar interaction difference matrix, which incorporates the partial charge of polar atoms in the ligand or protein. Similar to the hydrogen bond difference matrix, these interactions are expressed using simplified functions for the sake of the same reason. The values within these matrices are influenced by the square of the distance, a reflection of Coulomb potential. As with the hydrogen bond difference matrix, we utilize a smooth L1 loss function and mask other interactions to calculate the loss for each matrix, respectively. Here, denotes partial charge. Note that functional forms here are simplified to capture globular aspects; more detailed functional forms are used in the binding energy network instead.

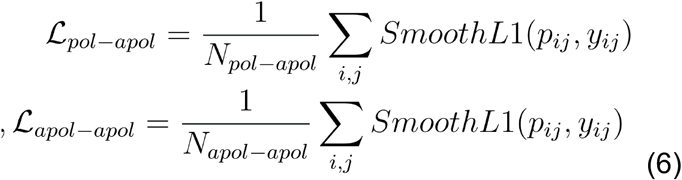

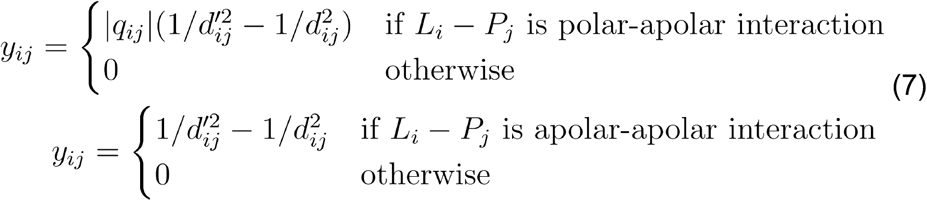

##### B.2 Weights of embedding vectors to predict lDDT

As described in Figure 3 (a), the embedding vectors are finally weighted to calculate complex and atom-wise lDDT. At this time, instead of simply using learnable weights, protein and ligand embeddings learned in the graph embedding module are used. Fixed learnable weights would remain unchanged after the learning process is complete, potentially limiting the model’s flexibility. However, by deriving weights from the learned embeddings, this allows the weights to adapt based on the context of the protein-ligand complex. The node embedding of the complex graph is projected into 5 dimensions using MLP and the weight is obtained by taking the softmax, which is used as the weight of the remaining 5 embeddings excluding the embedding obtained from the local atom graph. The local atom graph was fixed at a small weight of 0.1 to prevent overfitting due to increased dependence by learning very local information. This small weight ensures that the local atom graph acts as a minor correction term, which is sufficient for its intended role without greatly influencing the overall learning process.

##### B.3 EGNN blocks

The EGNN block utilized in the model architecture comprises multiple EGNN layers, as detailed in section 3.2.1. In the graph embedding module, we employ 4, 6, and 3 layers for the residue graph, atom graph, and local atom graph, respectively. The EGNN block used in the binding energy prediction network consists of 6 layers. The processes for node feature update and message passing in the EGNN layer are as follows.

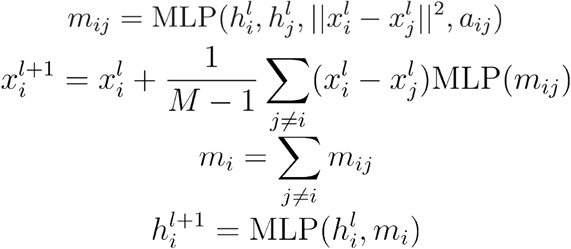

##### B.4 Theoretical background of the binding energy prediction network

In Poisson-Boltzmann (PB) or Generalized Born (GB) solvation models, the electrostatics interactions are coupled with the polar solvation so that the net electrostatic interaction strength varies on the environment. On the other hand, the Rosetta energy function uses a separate solvation term that takes into account of both polar and non-polar solvation effects together (non-polar solvation is usually counted by surface area models in PB or GB), while electrostatic screening effect is dampened by an environment-independent distance-dependent dielectric model. Because the network is aimed to reproduce individual terms in the Rosetta energy function, the theoretical implication of each term follows that of the Rosetta energy function but the fitting functions are brought from GB model. For instance, the solvation term in the network aims to fit GB self-solvation plus the surface area term. The electrostatic term in the network covers both original Coulombic term plus the polar pairwise term in GB.

##### B.5 Hyperparameters

The hyperparameters used in this paper are described in **Table S3**. Some parameters were selected empirically, but the limited computational resources prevented testing all possible combinations. Consequently, a more optimal combination may exist that was not identified. The protein-ligand complex is represented as a heterogeneous graph, with edges connecting nodes within a certain distance based on Euclidean distance. Additionally, we experimented with connecting the Top N nearest nodes, but this approach yielded suboptimal performance compared to distance-based connections. Therefore, the distance-based connection is selected.

##### B.6 Computational cost for training and inferencing

The number of learnable parameters in the Native-likeness prediction network and the Binding energy prediction network are 123,807 and 51,178, respectively. These models were trained on one 48GB A6000 GPU for about 100 epochs and each model was trained separately. Regarding the inference time, for 10 protein-ligand complex datasets, each containing 102 decoys (i.e., approximately 1,000 data points in total), the runtime on the CPU is about 125 seconds using 20 threads, while the runtime on the GPU is 63 seconds.

#### C Additional results

##### C.1 Docking with generated ligand conformations

**Table S4** shows the performance of model docking and rescoring results with ligand conformations generated by CORINA (v.5.0.0) on the PoseBusters model-docking benchmark set. Notably, the generated ligands affect the performance. However, as described above, the definition of success as complex lDDT over 0.9 is a stricter cutoff than the Ligand RMSD 2.0 criterion, making it more susceptible to conformational errors in the generated ligands. Therefore, we examined the success rate using 0.85 as the criterion and found that the Top1 success rate increased approximately twofold. Despite the sampling result not being optimal, DENOISer still increases the success rate in this scenario.

##### C.2 Fewer learnable parameters with physical inductive bias

The **Figure S5** illustrates the number of parameters utilized in the binding energy prediction network. Among all model parameters, the vdW, Electrostatic, and Solvation modules constitute a minimal proportion, with the majority being allocated to the EGNN block responsible for extracting atom embeddings. This approach, which incorporates physics-based inductive biases into the model architecture, allows for efficient achievement of high performance with a relatively small number of parameters.

#### D Limitations

##### D.1 Effects of pLDDT as an input feature

In the current model, plDDT is entered as an input feature, but it is not thought to play a significant role. Structural differences at the binding site critically impact the success rate in model docking. If such information could be used to assess the native-likeness of model docking, it would provide valuable insights for estimating the likeness. However, applying this information is challenging when the actual structure is unknown in the model docking scenario. Consequently, AF2’s plDDT can be employed as alternative information, as it reflects the confidence level of the local protein structure

We observed a correlation between the plDDT of the binding site and the complex lDDT, prompting us to include this information at the input feature level. Nonetheless, we believe there are potential methods to better leverage this information, enabling the model to respond more sensitively to plDDT. Exploring such methods could be conducted for future works.

##### D.2 Explicit solvation effect

In some structures of the protein-ligand complex, water molecules play a critical role in mediating interactions, such as forming hydrogen-bond bridges between the protein and ligand. In our current model, the effects of water molecules are not explicitly incorporated, relying instead on the assumption that the model will implicitly learn these interactions. However, explicitly including water molecules in the model could enhance the accuracy and specificity of the information provided to the model. Numerous methodologies exist for explicitly considering water molecules ^35,36^, yet these are not implemented in our current approach. Future research will involve integrating these methodologies to assess their impact on model performance. This analysis will provide insights into how explicitly accounting for water molecules may improve the prediction of protein-ligand interactions and overall model accuracy.

## Notes

### Competing Interest Statement

The authors have declared no competing interest.

## References

(1) Li, Y.; Li, L.; Wang, S.; Tang, X. EQUIBIND: A Geometric Deep Learning-Based Protein-Ligand Binding Prediction Method. Drug Discov. Ther. 2023, 17 (5), 363–364.

(2) Pei, Q.; Gao, K.; Wu, L.; Zhu, J.; Xia, Y.; Xie, S.; Qin, T.; He, K.; Liu, T.-Y.; Yan, R. FABind: Fast and Accurate Protein-Ligand Binding, 2023. http://arxiv.org/abs/2310.06763 (accessed 2024-06-23).

(3) Sutherland, J. J.; Nandigam, R. K.; Erickson, J. A.; Vieth, M. Lessons in Molecular Recognition. 2. Assessing and Improving Cross-Docking Accuracy. J. Chem. Inf. Model. 2007, 47 (6), 2293–2302.

(4) Lee, S.; Kim, S.; Lee, G. R.; Kwon, S.; Woo, H.; Seok, C.; Park, H. Evaluating GPCR Modeling and Docking Strategies in the Era of Deep Learning-Based Protein Structure Prediction. Comput. Struct. Biotechnol. J. 2023, 21, 158–167.

(5) Tunyasuvunakool, K.; Adler, J.; Wu, Z.; Green, T.; Zielinski, M.; Žídek, A.; Bridgland, A.; Cowie, A.; Meyer, C.; Laydon, A.; Velankar, S.; Kleywegt, G. J.; Bateman, A.; Evans, R.; Pritzel, A.; Figurnov, M.; Ronneberger, O.; Bates, R.; Kohl, S. A. A.; Potapenko, A.; Ballard, A. J.; Romera-Paredes, B.; Nikolov, S.; Jain, R.; Clancy, E.; Reiman, D.; Petersen, S.; Senior, A. W.; Kavukcuoglu, K.; Birney, E.; Kohli, P.; Jumper, J.; Hassabis, D. Highly Accurate Protein Structure Prediction for the Human Proteome. Nature 2021, 596 (7873), 590–596.

(6) Zheng, L.; Fan, J.; Mu, Y. OnionNet: A Multiple-Layer Intermolecular-Contact-Based Convolutional Neural Network for Protein-Ligand Binding Affinity Prediction. ACS Omega 2019, 4 (14), 15956–15965.

(7) Nguyen, D. D.; Wei, G.-W. AGL-Score: Algebraic Graph Learning Score for Protein–Ligand Binding Scoring, Ranking, Docking, and Screening. J. Chem. Inf. Model. 2019. 10.1021/acs.jcim.9b00334.

(8) Ragoza, M.; Hochuli, J.; Idrobo, E.; Sunseri, J.; Koes, D. R. Protein–Ligand Scoring with Convolutional Neural Networks. 2017. 10.1021/acs.jcim.6b00740.

(9) Durrant, J. D.; Andrew McCammon, J. NNScore: A Neural-Network-Based Scoring Function for the Characterization of Protein™Ligand Complexes. 2010. 10.1021/ci100244v.

(10) Durrant, J. D.; McCammon, J. A. NNScore 2.0: A Neural-Network Receptor-Ligand Scoring Function. J. Chem. Inf. Model. 2011, 51 (11), 2897–2903.

(11) Wang, R.; Fang, X.; Lu, Y.; Wang, S. The PDBbind Database: Collection of Binding Affinities for Protein-Ligand Complexes with Known Three-Dimensional Structures. J. Med. Chem. 2004, 47 (12), 2977–2980.

(12) Ramachandran, S.; Kota, P.; Ding, F.; Dokholyan, N. V. Automated Minimization of Steric Clashes in Protein Structures. Proteins 2011, 79 (1), 261–270.

(13) Jumper, J.; Evans, R.; Pritzel, A.; Green, T.; Figurnov, M.; Ronneberger, O.; Tunyasuvunakool, K.; Bates, R.; Žídek, A.; Potapenko, A.; Bridgland, A.; Meyer, C.; Kohl, S. A. A.; Ballard, A. J.; Cowie, A.; Romera-Paredes, B.; Nikolov, S.; Jain, R.; Adler, J.; Back, T.; Petersen, S.; Reiman, D.; Clancy, E.; Zielinski, M.; Steinegger, M.; Pacholska, M.; Berghammer, T.; Bodenstein, S.; Silver, D.; Vinyals, O.; Senior, A. W.; Kavukcuoglu, K.; Kohli, P.; Hassabis, D. Highly Accurate Protein Structure Prediction with AlphaFold. Nature 2021, 596 (7873), 583–589.

(14) Park, H.; Zhou, G.; Baek, M.; Baker, D.; DiMaio, F. Force Field Optimization Guided by Small Molecule Crystal Lattice Data Enables Consistent Sub-Angstrom Protein-Ligand Docking. J. Chem. Theory Comput. 2021, 17 (3), 2000–2010.

(15) Satorras, V. G.; Hoogeboom, E.; Welling, M. E(n) Equivariant Graph Neural Networks, 2021. http://arxiv.org/abs/2102.09844 (accessed 2024-06-23).

(16) Mariani, V.; Kiefer, F.; Schmidt, T.; Haas, J.; Schwede, T. Assessment of Template Based Protein Structure Predictions in CASP9. Proteins 2011, 79 Suppl 10, 37–58.

(17) Tyka, M. D.; Keedy, D. A.; André, I.; Dimaio, F.; Song, Y.; Richardson, D. C.; Richardson, J. S.; Baker, D. Alternate States of Proteins Revealed by Detailed Energy Landscape Mapping. J. Mol. Biol. 2011, 405 (2), 607–618.

(18) Buttenschoen, M.; Morris, G. M.; Deane, C. M. PoseBusters: AI-Based Docking Methods Fail to Generate Physically Valid Poses or Generalise to Novel Sequences, 2023. http://arxiv.org/abs/2308.05777 (accessed 2024-06-23).

(19) Aggarwal, R.; Gupta, A.; Priyakumar, U. D. APObind: A Dataset of Ligand Unbound Protein Conformations for Machine Learning Applications in De Novo Drug Design, 2021. http://arxiv.org/abs/2108.09926 (accessed 2024-06-23).

(20) Su, M.; Yang, Q.; Du, Y.; Feng, G.; Liu, Z.; Li, Y.; Wang, R. Comparative Assessment of Scoring Functions: The CASF-2016 Update. J. Chem. Inf. Model. 2018. 10.1021/acs.jcim.8b00545.

(21) Liu, Z.; Su, M.; Han, L.; Liu, J.; Yang, Q.; Li, Y.; Wang, R. Forging the Basis for Developing Protein-Ligand Interaction Scoring Functions. Acc. Chem. Res. 2017, 50 (2), 302–309.

(22) Trott, O.; Olson, A. J. AutoDock Vina: Improving the Speed and Accuracy of Docking with a New Scoring Function, Efficient Optimization, and Multithreading. J. Comput. Chem. 2010, 31 (2), 455–461.

(23) McNutt, A. T.; Francoeur, P.; Aggarwal, R.; Masuda, T.; Meli, R.; Ragoza, M.; Sunseri, J.; Koes, D. R. GNINA 1.0: Molecular Docking with Deep Learning. J. Cheminform. 2021, 13 (1), 43.

(24) Corso, G.; Stärk, H.; Jing, B.; Barzilay, R.; Jaakkola, T. DiffDock: Diffusion Steps, Twists, and Turns for Molecular Docking, 2022. http://arxiv.org/abs/2210.01776 (accessed 2024-06-23).

(25) Shin, W.-H.; Seok, C. GalaxyDock: Protein-Ligand Docking with Flexible Protein Side-Chains. J. Chem. Inf. Model. 2012, 52 (12), 3225–3232.

(26) Kelley, B. P.; Brown, S. P.; Warren, G. L.; Muchmore, S. W. POSIT: Flexible Shape-Guided Docking For Pose Prediction. J. Chem. Inf. Model. 2015, 55 (8), 1771–1780.

(27) Francoeur, P. G.; Masuda, T.; Sunseri, J.; Jia, A.; Iovanisci, R. B.; Snyder, I.; Koes, D. R. Three-Dimensional Convolutional Neural Networks and a Cross-Docked Data Set for Structure-Based Drug Design. J. Chem. Inf. Model. 2020. 10.1021/acs.jcim.0c00411.

(28) Kwon, S.; Seok, C. CSAlign and CSAlign-Dock: Structure Alignment of Ligands Considering Full Flexibility and Application to Protein–ligand Docking. Comput. Struct. Biotechnol. J. 2023, 21, 1–10.

(29) Krishna, R.; Wang, J.; Ahern, W.; Sturmfels, P.; Venkatesh, P.; Kalvet, I.; Lee, G. R.; Morey-Burrows, F. S.; Anishchenko, I.; Humphreys, I. R.; McHugh, R.; Vafeados, D.; Li, X.; Sutherland, G. A.; Hitchcock, A.; Hunter, C. N.; Kang, A.; Brackenbrough, E.; Bera, A. K.; Baek, M.; DiMaio, F.; Baker, D. Generalized Biomolecular Modeling and Design with RoseTTAFold All-Atom. Science 2024, 384 (6693), eadl2528.

(30) Qiao, Z.; Nie, W.; Vahdat, A.; Miller, T. F.; Anandkumar, A. State-Specific Protein–ligand Complex Structure Prediction with a Multiscale Deep Generative Model. Nature Machine Intelligence 2024, 6 (2), 195–208.

(31) Abramson, J.; Adler, J.; Dunger, J.; Evans, R.; Green, T.; Pritzel, A.; Ronneberger, O.; Willmore, L.; Ballard, A. J.; Bambrick, J.; Bodenstein, S. W.; Evans, D. A.; Hung, C.-C.; O’Neill, M.; Reiman, D.; Tunyasuvunakool, K.; Wu, Z.; Žemgulytė, A.; Arvaniti, E.; Beattie, C.; Bertolli, O.; Bridgland, A.; Cherepanov, A.; Congreve, M.; Cowen-Rivers, A. I.; Cowie, A.; Figurnov, M.; Fuchs, F. B.; Gladman, H.; Jain, R.; Khan, Y. A.; Low, C. M. R.; Perlin, K.; Potapenko, A.; Savy, P.; Singh, S.; Stecula, A.; Thillaisundaram, A.; Tong, C.; Yakneen, S.; Zhong, E. D.; Zielinski, M.; Žídek, A.; Bapst, V.; Kohli, P.; Jaderberg, M.; Hassabis, D.; Jumper, J. M. Accurate Structure Prediction of Biomolecular Interactions with AlphaFold 3. Nature 2024, 630 (8016), 493–500.

(32) Bryant, P.; Kelkar, A.; Guljas, A.; Clementi, C.; Noé, F. Structure Prediction of Protein-Ligand Complexes from Sequence Information with Umol. Nat. Commun. 2024, 15 (1), 1–12.

(33) Li, W.; Godzik, A. Cd-Hit: A Fast Program for Clustering and Comparing Large Sets of Protein or Nucleotide Sequences. Bioinformatics 2006, 22 (13), 1658–1659.

(34) Alford, R. F.; Leaver-Fay, A.; Jeliazkov, J. R.; O’Meara, M. J.; DiMaio, F. P.; Park, H.; Shapovalov, M. V.; Renfrew, P. D.; Mulligan, V. K.; Kappel, K.; Labonte, J. W.; Pacella, M. S.; Bonneau, R.; Bradley, P.; Dunbrack, R. L., Jr; Das, R.; Baker, D.; Kuhlman, B.; Kortemme, T.; Gray, J. J. The Rosetta All-Atom Energy Function for Macromolecular Modeling and Design. J. Chem. Theory Comput. 2017, 13 (6), 3031–3048.

(35) Park, S.; Seok, C. GalaxyWater-CNN: Prediction of Water Positions on the Protein Structure by a 3D-Convolutional Neural Network. J. Chem. Inf. Model. 2022, 62 (13), 3157–3168.

(36) Sato, K.; Oide, M.; Nakasako, M. Prediction of Hydrophilic and Hydrophobic Hydration Structure of Protein by Neural Network Optimized Using Experimental Data. Sci. Rep. 2023, 13 (1), 1–15.

